# Optogenetic *lac* operon to control chemical and protein production in *Escherichia coli* with light

**DOI:** 10.1101/845453

**Authors:** Makoto A. Lalwani, Samantha S. Ip, Cesar Carrasco-Lopez, Evan M. Zhao, Hinako Kawabe, José L. Avalos

## Abstract

Control of the *lac* operon with IPTG has been used for decades to regulate gene expression in *E. coli* for countless applications, including metabolic engineering and recombinant protein production. However, optogenetics offers unique capabilities such as easy tunability, reversibility, dynamic induction strength, and spatial control that are difficult to obtain with chemical inducers. We developed an optogenetic *lac* operon in a series of circuits we call OptoLAC. With these circuits, we control gene expression from various IPTG-inducible promoters using only blue light. Applying them to metabolic engineering improves mevalonate and isobutanol production by 24% and 27% respectively, compared to IPTG induction, in light-controlled fermentations scalable to at least 2L bioreactors. Furthermore, OptoLAC circuits enable light control of recombinant protein production, reaching yields comparable to IPTG induction, but with enhanced tunability of expression and spatial control. OptoLAC circuits are potentially useful to confer light controls over other cell functions originally engineered to be IPTG-inducible.

## Introduction

The long history of *Escherichia coli* as a model organism for basic and applied research makes it a preferred host for many biotechnological applications, including metabolic engineering for chemical production and expression of recombinant proteins. The vast wealth of knowledge on the physiology, genetics, and metabolism of this bacterium, as well as the abundance of molecular tools for its genetic manipulation, have made *E. coli* an organism of choice across the spectrum of metabolic engineering, from proofs of principle^1^ to the development of large-scale industrial processes^2–5^. However, metabolic engineering in *E. coli* still faces important challenges, especially considering that each pathway presents distinct obstacles, which may be addressed with technological improvements.

A key challenge in metabolic engineering is the need for dynamic control^6^. This need commonly arises when the biosynthesis of the product of interest competes with cell growth, or when the product of interest, or its precursors, are toxic to the host organism, leading to growth inhibition and poor productivity. These challenges are usually addressed by dividing fermentations into a growth phase (in which the biosynthetic pathway of interest is repressed and metabolism is devoted to growing biomass) and a production phase (in which the pathway of interest is chemically induced, frequently at the expense of cell growth). However, pathway productivities can be greatly impacted by the timing, rates, and levels of induction, which can be difficult to control with chemical inducers. We recently showed that light can be an effective alternative to chemicals as an inducing agent for metabolic engineering in *S. cerevisiae*, enhancing the dynamic control over fermentations by adding tunability, reversibility, and independence from medium composition^7^. To achieve this, we devised optogenetic circuits that harness the regulon system that responds to galactose, which is frequently utilized to induce gene expression in yeast.

The most commonly used inducible system in *E. coli* is based on the *lac* operon^8^, more specifically its repressor LacI. Genes required for lactose utilization are naturally repressed in the absence of this sugar or in the presence of a preferred carbon source (i.e. glucose). This repression is mediated by LacI binding to *lac* operator (*lacO*) sites located upstream of genes involved in lactose metabolism. When glucose is consumed, lactose binding to LacI causes it to dissociate from *lacO* sites, thus allowing gene transcription. For decades, this system has been exploited for dynamic control of gene expression in *E. coli*, using *lacO* sites to recruit LacI to a variety of promoters, excluding catabolite activator protein (CAP)-binding sequences to prevent glucose repression, and using isopropyl β-D-1-thiogalactopyranoside (IPTG), a non-metabolizable allolactose mimetic, as an inducing agent^9^. These features of the *lac* operon provide an opportunity to develop optogenetic circuits to control gene expression in *E. coli* with light.

Several optogenetic systems have been developed in *E. coli*^10–15^. Among these, the photoreceptor signaling cascade encoded in the pDawn plasmid^15^ seemed that it would be particularly adept at harnessing the *lac* operon. The pDawn system is derived from a blue light-responsive photosensory histidine kinase YF1 and its cognate response transcriptional regulator FixJ from *Bradyrhizobium japonicum*. In the dark, the membrane-bound YF1 phosphorylates FixJ, which then activates the P_FixK2_ promoter to express the λ phage repressor cI. This repressor in turn prevents transcription from its cognate P_R_ promoter, which controls genes of interest. Conversely, blue light (∼470 nm) induces phosphatase activity in YF1^16^, thus preventing FixJ phosphorylation and cI expression. With this design, genes controlled by the P_R_ promoter are repressed in the dark by cI, and expressed in blue light by the absence of this repressor.

In this study, we report the development of an optogenetic *lac* operon in a series of circuits we call OptoLAC. We achieve this by controlling expression of the *lacI* repressor using the P_R_ promoter from the pDawn system. These optogenetic circuits can control promoters originally designed to be IPTG-inducible, making them tunable to intermediate levels of expression by using different light duty cycles. Moreover, our OptoLAC circuits can be readily used to replace IPTG with light as an inducing agent to control engineered metabolic pathways for chemical production in lab-scale bioreactors, reaching titers of mevalonate and isobutanol that are higher than those achieved with IPTG induction. In addition, we develop an OptoLAC circuit to control gene expression in *E.coli* strains for recombinant protein production. These new optogenetic circuits open the door to using light control for metabolic engineering and protein production in *E. coli*, as well as potentially for other research and biotechnological applications currently controlled by IPTG.

## Results

### Development of optogenetic *lac* operon circuits to control gene expression

To develop light controls for chemical and protein production in *E. coli*, we hijacked the lac repressor (LacI). We placed *lacI* under the cI-controlled P_R_ promoter of pDawn, such that genes for metabolic pathways or protein production are repressed in the light and induced in the dark (Figure 1a). To ensure control over total LacI levels, we adapted a *lacI*-deficient *E. coli* K-12 MG1655 strain^17^ (EMAL52, hereafter referred to as OptoMG), which we transformed with plasmids (pMAL288 and pMAL292, Supplementary Table 1) to control *lacI* expression with pDawn, and super-folder green fluorescent protein (GFP) expression with the IPTG-inducible promoter P_T5-lacO_, resulting in strain EMAL57 (Supplementary Table 2). Instead of inducing GFP expression in EMAL57 with IPTG, we grew it in either light or dark conditions for 8 hours. However, there is only a 1.5-fold increase in GFP expression when EMAL57 is grown in the dark relative to being grown in the light. Furthermore, GFP expression under either light condition is less than 1% of the levels achieved with the strong constitutive promoter P_BBa_J23100_ (P_BBA_), which also serves as a control to show that GFP does not significantly bleach under our light conditions (Supplementary Figure 1a). These results indicate that pDawn control of *lacI* expression requires further engineering to achieve robust optogenetic control of gene expression.

**Figure 1.**
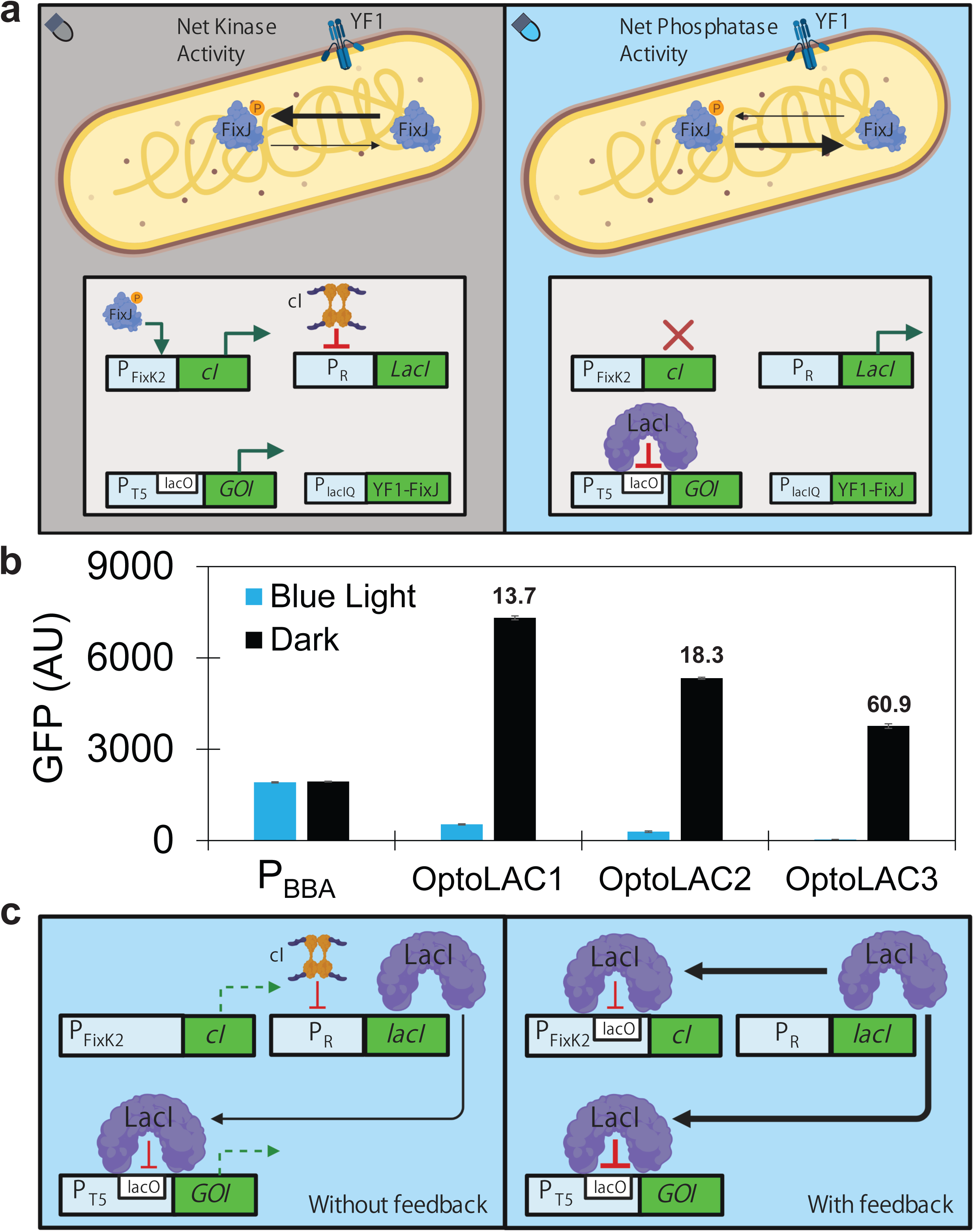
OptoLAC Circuit schematic. **(a)** OptoLAC circuits use the pDawn optogenetic system to control *lacI* expression. In the dark, the histidine kinase YF1 phosphorylates its response regulator FixJ, allowing it to activate gene expression from its cognate P_FixK2_ promoter (left panel). This P_FixK2_ promoter controls expression of the λ phage repressor cI, which represses the P_R_ promoter. Thus, by controlling *lacI* expression with the P_R_ promoter, cI-mediated repression of *lacI* allows *lacO*-containing promoters to express genes of interest (*GOI*) in the dark. On the other hand, blue light inhibits YF1, leaving the unphosphorylated FixJ unable to activate P_FixK2_ and express cI (right panel). In the absence of cI, the P_R_ promoter is able to express LacI, which then binds *lacO* sites. Therefore genes of interest (*GOI*) controlled by *lacO*-containing promoters are repressed in blue light. **(b)** GFP expression from the strong constitutive promoter P_BBA_ (EMAL231), or from P_T5-lacO_ controlled by OptoLAC1 (EMAL68), OptoLAC2 (EMAL69), OptoLAC3 (EMAL230), or, in blue light (blue) or darkness (black). The fold of induction of each OptoLAC circuit is shown on top of their corresponding graph. All data shown as median values of 10,000 single-cell flow cytometry events; error bars represent one standard deviation of replicates exposed to the same conditions (n = 3 biologically independent 150 µL samples). All experiments were repeated at least three times. **(c)** Schematic showing insertion of lacO site downstream of P_FixK2_ to reduce leaky cI transcription and promote LacI expression, thereby reducing leaky expression under blue light. Without feedback (left panel), leaky expression of cI under blue light (dashed green arrow) causes repression of *lacI* transcription, resulting in leaky transcription of genes of interest (dashed green arrow). With feedback (right panel), tight repression of cI transcription under blue light enables uninhibited transcription of *lacI*, which results in increased LacI protein levels (bold dark arrows) and tight repression of genes of interest (bold red line).

The poor induction and overall low levels of GFP expression in EMAL57 suggest that the levels of LacI in this strain are too high regardless of cI activity. To address this, we reduced the half-life of LacI by fusing to its C-terminus an ssRA degradation tag^18,19^ of three possible sequences differing only in the final three residues (LAA, AAV, or ASV), which impact tag strength^20,21^. Two of the three resulting strains (EMAL68, EMAL69, and EMAL71) show darkness-induced GFP expression levels that exceed constitutive expression from P_BBA_ (Supplementary Figure 1b). Using the strongest ssRA-LAA tag results in the highest maximal GFP expression in the dark, consistent with it having the most destabilizing effect on LacI. However, the ssRA-AAV tag offers tighter control of gene expression under blue light, likely due to higher LacI-mediated repression. Therefore, we continued our study using LacI fusions to ssRA-LAA and ssRA-AAV because they are both effective at controlling gene expression with light, naming them OptoLAC1 (inducible to 3.78-fold higher expression than P_BBA_, with 13.7-fold induction) and OptoLAC2 (inducible to 2.75-fold higher expression than P_BBA_, with 18.3-fold induction), respectively (Figure 1b). These results reflect the need to carefully manage the levels of LacI in order to achieve robust optogenetic control over the *lac* operon gene expression system.

While fusing LacI to degradation tags is effective at improving the levels of induction of GFP expression in the dark, the reduced stability of LacI results in relatively leaky expression of GFP in the light with both OptoLAC1 and OptoLAC2 (Figure 1b). To reduce the leaky expression of OptoLAC2 in the light and improve its fold of induction, we built a circuit with positive feedback to increase the repression of *lacI* in the light. We introduced a *lacO* operator sequence downstream of the core sequence of the P_FixK2_ promoter that drives the expression of the cI repressor (Supplementary Sequence 2). With this design, LacI represses transcription of both the gene of interest (e.g. GFP) and the cI repressor that controls its own expression, thus increasing transcription of *lacI* in blue light and resulting in a positive feedback loop that tightens circuit repression (Figure 1c). This new circuit, which we named OptoLAC3, shows lower GFP expression (EMAL230) in blue light than OptoLAC1 or OptoLAC2 (Figure 1c). A tradeoff of reducing leakiness by inserting a *lacO* operator sequence downstream of P_FixK2_ is that it weakens this promoter (Supplementary Figure 1c), which lowers the cI-mediated repression of *lacI* in the dark, and thus the maximal GFP expression achieved by OptoLAC3. Nevertheless, OptoLAC3 achieves a 60.9-fold induction of gene expression in the dark, which is the highest of the OptoLAC circuits. OptoLAC1, 2, and 3 represent the first suite of optogenetic circuits that harness the *lac* operon for gene expression in *E. coli*, which enable the use of light to control processes originally designed to be IPTG inducible.

### Characterization of OptoLAC circuits

To test the tunability of our OptoLAC circuits, we grew strains containing OptoLAC1 (EMAL68) or OptoLAC3 (EMAL230) under different light doses, measured in time fractions of light exposure. We found that OptoLAC1 requires only 10% of light exposure (100 seconds of light per 1000 seconds) to reach 99% of maximum repression under full light (Figure 2a). In contrast, OptoLAC3 is significantly more sensitive, achieving full repression under only 1% of light exposure. The differences in light sensitivities between OptoLAC circuits may be useful for different applications. For example, it may be easier to maintain intermediate levels of expression in the less sensitive OptoLAC1 circuit, while OptoLAC3 may be better suited to achieve full gene repression in conditions where light penetration may be limited.

**Figure 2.**
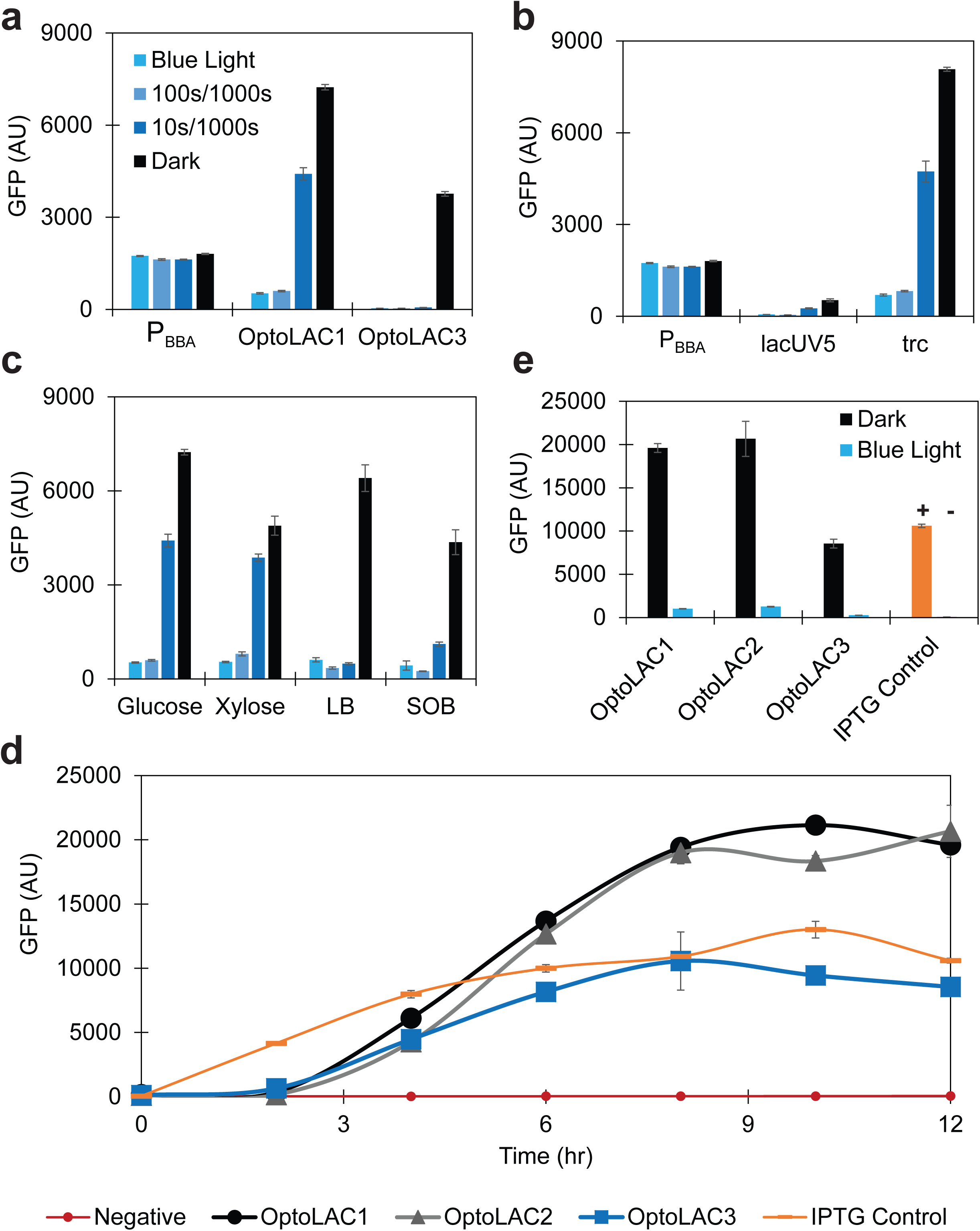
Circuit characterization. **(a)** GFP expression from P_BBA_ (EMAL231) or from P_T5-lacO_ controlled by OptoLAC1 (EMAL68) or OptoLAC3 (EMAL230), under different doses of blue light: full light (100% light), 100 sec ON/900 sec OFF (10% light), 10 sec ON/990 sec OFF (1% light), and full darkness (0% light). **(b)** GFP expression from P_BBA_ (EMAL231) or controlled by OptoLAC1 from other LacI-repressed promoters: P_lacUV5_ (EMAL267) and P_trc_ (EMAL268). **(c)** GFP expression from P_T5-lacO_ controlled by OptoLAC1 (EMAL68) in M9 + 2% glucose, M9 + 2% xylose, LB, and SOB. **(d)** Time course of GFP expression from P_T5-lacO_ controlled by OptoLAC1 (EMAL68), OptoLAC2 (EMAL69), OptoLAC3 (EMAL230), or IPTG (EMAL77) compared to a negative control lacking GFP (EMAL229), which also shows the low level of autofluorescence. Strains were grown under light until exponential phase, then switched to full darkness (or EMAL77 induced with 1 mM IPTG) at t = 0. **(e)** GFP expression in each culture at the 12 hour mark in the time course shown in (d), including uninduced controls where cultures were kept in blue light, or no IPTG was added. All data shown as median values of 10,000 single-cell flow cytometry events; error bars represent one standard deviation of replicates exposed to the same conditions (n = 3 biologically independent 150 µL or 1 mL samples). Experiments were repeated at least 3 times.

The original designs of OptoLAC circuits employ the IPTG-inducible P_T5-lacO_ promoter to express GFP. However, other promoters have been developed with *lacO* sequences to make them IPTG-inducible. To test whether our circuits can be used to control other *lacO*-containing promoters with light, we used OptoLAC1 to control GFP expression from P_lacUV5_^22^ and P_trc_^23^. We found that both promoters show light-tunable repression with varying degrees of maximal gene expression in the dark according to the core promoter strength (Figure 2b). This is an important finding, as it demonstrates that our OptoLAC circuits can be used to control different promoters originally designed to be inducible with IPTG, which facilitates the retrofitting of many existing systems with light controls.

Inducible promoters are frequently used for dynamic control in metabolic engineering for the production of fuels and chemicals from renewable sources, such as glucose or xylose. Gene induction is also commonly used in *E. coli* for the production of recombinant proteins, which may require different cultivation media, including the rich lysogeny broth (LB) and super optimal broth (SOB). Therefore, we tested the performance of OptoLAC1 in glucose and xylose, using M9 minimal salts media, as well as LB and SOB. Using OptoLAC1, it is possible to control gene expression with light in all media tested (Figure 2c). However, we observed some interesting differences; for example, OptoLAC1 seems to be more light-sensitive in rich media than in minimal media. In addition, while the light-sensitivity in glucose or xylose is similar, the maximal gene expression in the dark is greater in glucose. Our OptoLAC circuits are not only effective at controlling gene expression temporally in liquid media, but also spatially in solid media. Using OptoLAC1, we are able to selectively activate GFP expression in sections of an LB agar plate that are shielded from blue light (Supplementary Figure 2). These results raised the prospect of using our OptoLAC circuits for metabolic engineering and protein production.

Before testing our optogenetic *lac* operon circuits in biotechnological applications, we sought to compare them with IPTG, the current gold standard method to induce genes controlled by the *lac* operon in metabolic engineering and protein production. Given that the LacI degradation process involved in our optogenetic circuits is inherently slower than the fast-acting allosteric inhibition of LacI by IPTG, we reasoned that the most appropriate way to compare these systems would be in a time course. We grew in blue light strains carrying each OptoLAC circuit; when they reached exponential growth phase, we induced GFP expression by moving the cultures into the dark. As a control, we also used IPTG to induce GFP expression from a plasmid of equivalent copy number to our OptoLAC circuits but containing constitutively expressed *lacI* (pET28a).

Expression of GFP from OptoLAC circuits is detectable 2-4 hours after switching the light off, whereas, as expected, IPTG induction responds more rapidly with clear GFP expression after 2 hours of induction. However, it still takes 10 hours to reach full induction of GFP with IPTG, which is comparable to the time it takes for our optogenetic circuits to reach full induction (8-12 hours). After ∼4-5 hours of induction, the levels of GFP expression obtained with OptoLAC1 and OptoLAC2 are approximately the same as what IPTG induction achieves, while OptoLAC3 matches IPTG levels of induction after 8 hours. At their maximum levels of gene expression (approximately 10 hours after induction), the levels achieved with OptoLAC1 and OptoLAC2 are higher than those achieved by IPTG induction by 63% and 41%, respectively. Importantly, exposing the cultures containing OptoLAC circuits to blue light for the entire time course keeps GFP expression low (Figure 2e). These results show that despite their initial delay in induction, our optogenetic circuits eventually catch up and even exceed the induction levels achieved with IPTG.

### OptoLAC circuits for dynamic control of chemical production

The favorable performance of our OptoLAC circuits to control GFP expression compared to IPTG induction encouraged us to test them in light-controlled bacterial fermentations for chemical production. We first tested our optogenetic circuits to control the biosynthesis of mevalonate, an important terpenoid precursor previously produced in *E. coli* (Figure 3a)^24–26^. We adapted to our optogenetic platform a previously described IPTG-inducible plasmid^24^ containing the three genes to produce mevalonate from acetyl-CoA (*atoB*, *HMGS*, and *tHMGR*), resulting in plasmid pMAL487 (see Methods). Transforming our OptoMG chassis strain with pMAL487 and a plasmid containing each one of our OptoLAC circuits, we obtained strains in which the mevalonate biosynthetic pathways is under light control with OptoLAC1 (EMAL208), OptoLAC2 (EMAL209), or OptoLAC3 (EMAL235). As a control for IPTG-induced mevalonate production, we transformed OptoMG with pMAL487 and pET28a (for constitutive *lacI* expression) to make EMAL135.

**Figure 3.**
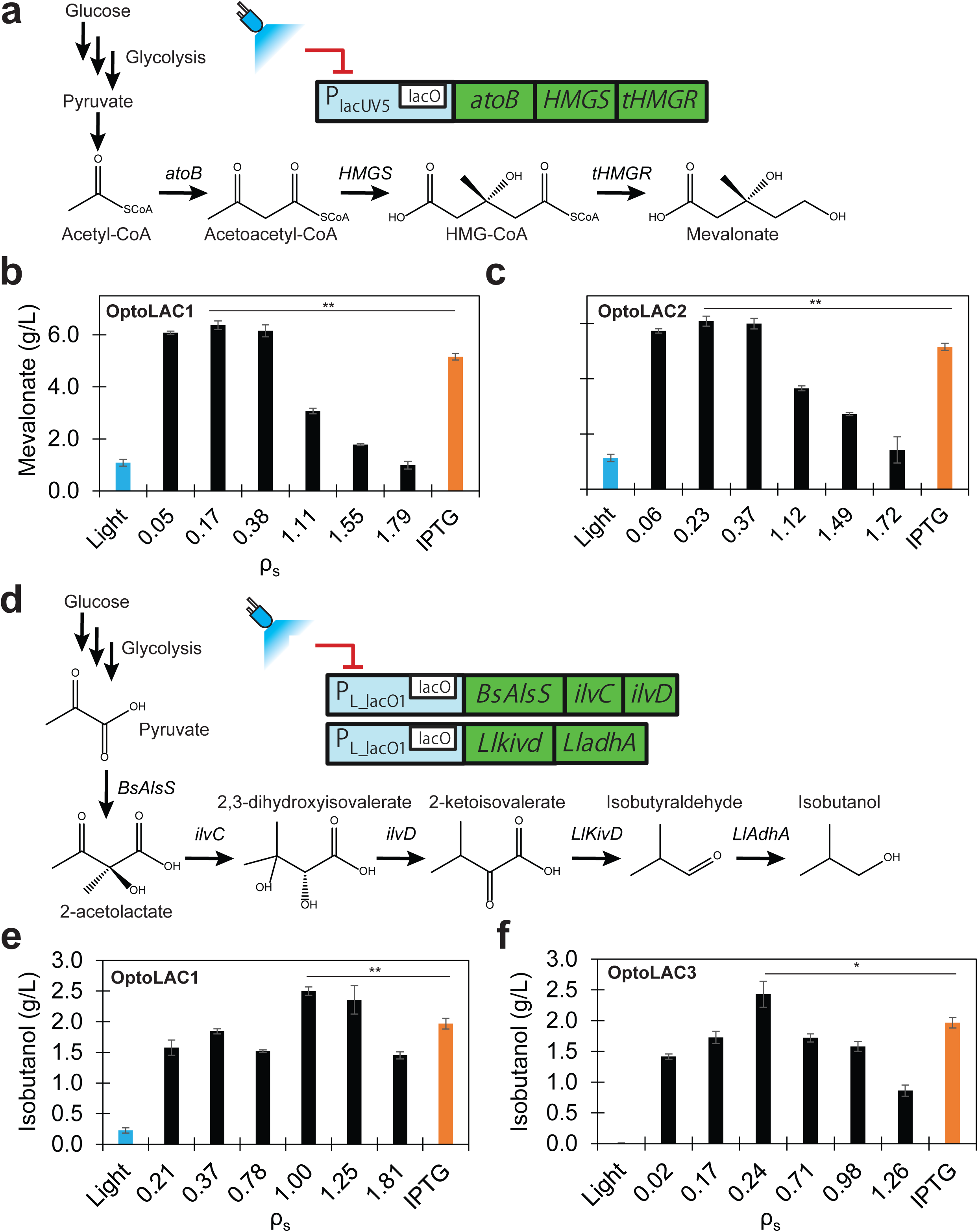
Chemical production. **(a)** Optogenetic control of the mevalonate biosynthetic pathway, composed of *atoB*, *HGMS*, and *tHMGR,* expressed from P_lacUV5_. **(b,c)** Mevalonate production controlled with **(b)** OptoLAC1 (EMAL208), or **(c)** OptoLAC2 (EMAL209) when switching cultures from light to dark at different cell densities (ρ_s_ values, black) or uninduced (blue); as well fermentations induced with IPTG (EMAL135) at optimal cell density (OD_600_ = 0.06, orange). **(d)** Optogenetic control of the isobutanol biosynthetic pathway, composed of *BsAlsS, ilvC, ilvD, LlKivD*, and *LlAdhaA*, expressed from P_L-LacO1_. **(e,f)** Isobutanol production controlled with **(e)** OptoLAC1 (EMAL199), or **(f)** OptoLAC3 (EMAL238) for different ρ_s_ values (black) or uninduced (blue); as well as fermentations induced by IPTG (EMAL201) at optimal cell density (OD_600_ = 1.4, orange). All data are shown as mean values; error bars represent the standard deviation of at least three biologically independent 1-mL sample replicates exposed to the same conditions. \**P* < 0.05, \*\**P* < 0.01. All experiments were repeated at least three times.

Light-controlled fermentations were grown in blue light to different OD_600_ before inducing the mevalonate pathway by switching the light off (see Methods). The control strain was also induced at different OD_600_ values, but with 1mM IPTG. After 72 hours of fermentations, we found that the OD_600_ at which we induce the culture by switching from light to dark conditions (ρ_s_) has a significant impact on the final mevalonate titers (Figure 3b,c). OptoLAC1 and OptoLAC2 achieve maximum mevalonate titers of 6.4 ± 0.2 g/L and 6.1 ± 0.2 g/L, respectively, at similar ρ_s_ values (OD_600_ ∼ 0.17-0.23), which exceed the titer obtained with IPTG induction at its optimal OD_600_ of induction by as much as 24% (Figure 3b,c and Supplementary Figure 3a). These results demonstrate that by optogenetically controlling the *lac* operon, it is possible to replace IPTG with light as an inducing agent in *E. coli* fermentations for chemical production.

To test whether our optogenetic circuits can be more generally applied to other metabolic pathways, we used them to control a second biosynthetic pathway; this time for the production of isobutanol, an advanced biofuel that presents different challenges to produce than mevalonate, including its higher cellular toxicity. Again, we adapted a previously described plasmid^27,28^ to our optogenetic platform (see Methods), resulting in plasmid pMAL534, which contains an IPTG-inducible isobutanol pathway (Figure 3d) comprised of five enzymes (*BsAlsS, ilvC, ilvD, LlkivD, LladhA*). We used pMAL534 to produce strains with the isobutanol pathway controlled by OptoLAC1 (EMAL199), OptoLAC2 (EMAL200), or OptoLAC3 (EMAL239), as well as a control strain inducible only with IPTG (EMAL201). As described above for mevalonate production, we conducted fermentations to test our circuits for isobutanol production using different ρ_s_ values (the OD_600_ at which we induce the isobutanol pathway with darkness). We found that for isobutanol production, OptoLAC1 and OptoLAC3 are the best performing circuits (Figure 3e,f). Unlike the case for light-controlled mevalonate fermentations, the optimal ρ_s_ values for isobutanol production differ significantly between OptoLAC1 (ρ_s_ = 1.0) and OptoLAC3 (ρ_s_ = 0.24). Furthermore, the darkness-induced isobutanol fermentations controlled from either OptoLAC circuit at their optimal ρ_s_ value achieve as much as 27% higher isobutanol titers (2.5 ± 0.1 g/L and 2.4 ± 0.2 g/L with OptoLAC1 and OptoLAC3, respectively) than the same plasmid induced by IPTG at its optimal OD_600_ of induction (2.0 ± 0.1 g/L, OD_600_ = 1.4, Supplementary Figure 3b). These results confirm that our OptoLAC circuits can be applied to control different metabolic pathways with light, improving the titers obtained with IPTG induction.

### Scaleup of light-controlled fermentation in bioreactors

OptoLAC circuits are intentionally designed to induce gene expression in the dark to circumvent the potential challenge of limited light penetration in high cell-density fermentations. To test the efficacy of this approach, we used EMAL208 (containing OptoLAC1) to produce mevalonate in a light-controlled 2-L bioreactor. We grew EMAL208 in M9 minimal salts medium and 5% glucose, with temperature, pH, and dissolved oxygen controls, as well as blue light panels illuminating approximately 77% of the fermentation bulk surface (Figure 4a). When the culture reached the previously determined optimal ρ_s_ (OD_600_ = 0.17), we induced the mevalonate pathway by turning off the LED panels and covering the bioreactor with aluminum foil and dark cloth, and started collecting samples for analysis throughout 52 hours of the fermentation. We found that for the first 3 hours after induction, mevalonate production is undetectable, demonstrating that blue light effectively represses the mevalonate pathway during the growth phase of the fermentation (Figure 4b). By 6 hours after induction, we see a rapid increase in mevalonate production, reaching a maximum titer of 6.1 g/L after 48 hours of induction, which replicates what we observe in small-scale batch fermentations. These results show that light is an effective agent to control *E. coli* fermentations in lab-scale bioreactors of up to at least 2-L.

**Figure 4.**
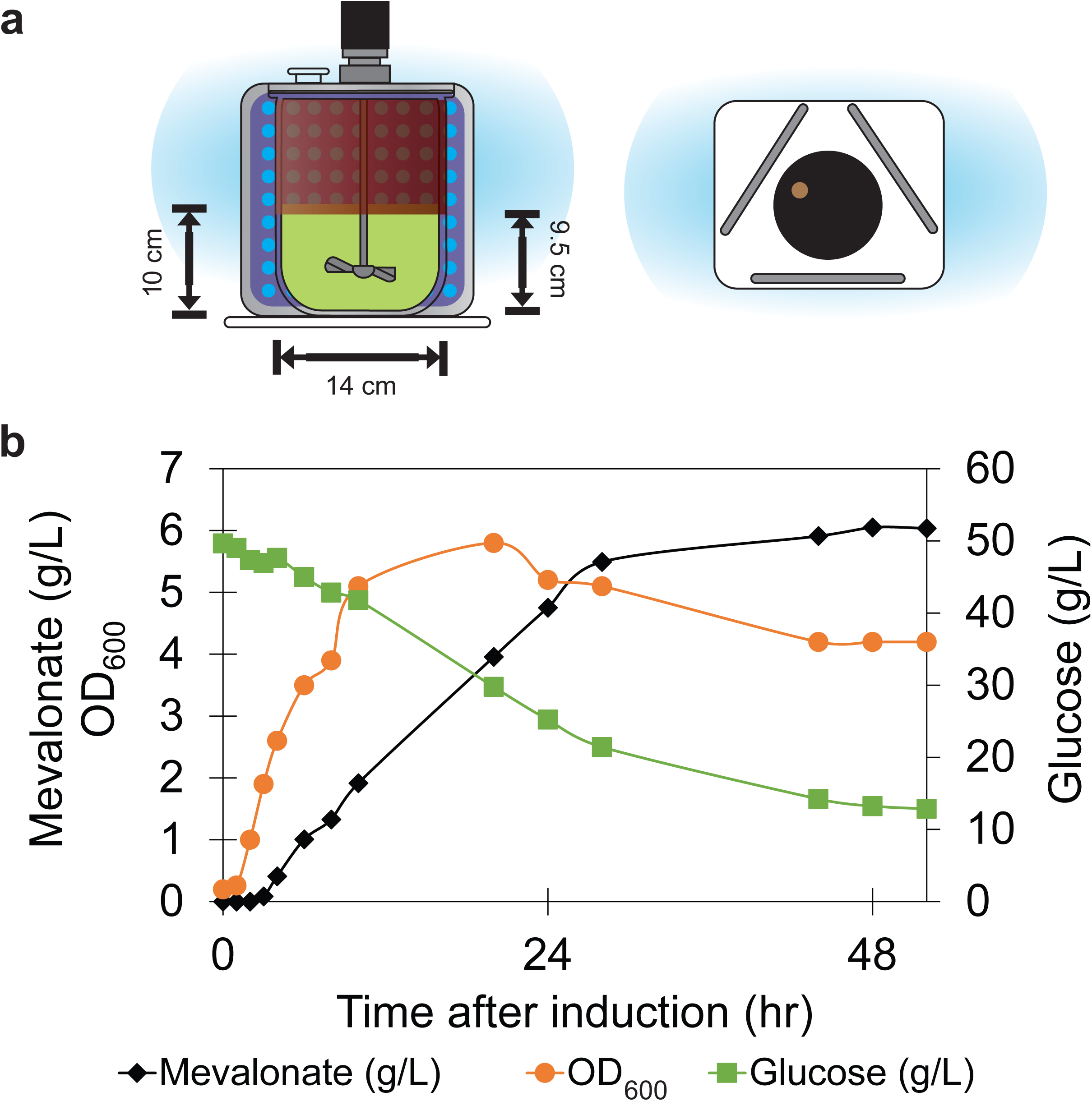
Scale-up of mevalonate production. a) Side- and top-view of a light-controlled 2-L bioreactor for mevalonate production, with three light panels arranged triangularly ∼20 cm from the glass vessel wall such that the light intensity at the vessel surface was between 80-110 µmol/m^2^/s. Elevating the heating blanket (red) allows exposure of ∼76% of the fermentation bulk surface to light. b) Light-controlled mevalonate production in a 2-L bioreactor. Measurements of mevalonate (black), glucose (purple), and OD_600_ (orange) taken after induction and throughout 52 hours of fermentation. EMAL208 was inoculated into a working volume of 1 L of M9 + 5% glucose and grown for 4 hours under blue light until reaching OD_600_ = 0.17, at which point the light panels were removed and the reactor was covered in aluminum foil and black cloth to protect it from ambient light. Samples were periodically taken and analyzed with HPLC.

### Optogenetic control of recombinant protein production

Our OptoLAC circuits are designed to control, in principle, any LacI-regulated (IPTG-inducible) promoter, so we set out to test them for light control of recombinant protein production, another common application of IPTG induction in *E. coli*. However, the strains used for protein production (usually B strain-derived) are different from those commonly used in metabolic engineering (usually K-12-derived). Therefore, we adapted BL21 (DE3), a workhorse strain for protein expression, to work with our optogenetic circuits. This involved deleting the two endogenous copies of *lacI* (see Methods) to produce EMAL255, which we call OptoBL. In addition, two commonly used protein expression vectors, pET and pCri, share the same origin of replication and selection marker as our plasmids carrying OptoLAC circuits (pMAL292, pMAL296, or pMAL630). Therefore, we reconstructed OptoLAC1 into a pACYC-derived^29^ backbone, resulting in OptoLAC1B (pMAL658), which is compatible with pET and pCri and is specifically designed to control protein production optogenetically in OptoBL (Supplementary Table 1). Finally, we adapted pCri-8b, which was originally designed to assay IPTG-induced yellow fluorescent protein (YFP) production from the P_T7_ promoter^30^, by removing its constitutive copy of *lacI* ^31^ (see Methods), resulting in pC85 (Supplementary Table 1).

With these tools in hand, we produced EMAL284 (Supplementary Table 2), which can be induced by darkness to produce high levels of YFP. We grew EMAL284 in blue light in LB medium. As a control, we grew an IPTG-inducible strain carrying pCri-8b (EMAL283) under similar conditions, but in ambient light. When the cultures reached exponential phase (OD_600_ = 0.5), we induced them by either turning off the light and covering with aluminum foil (EMAL284) or by adding 1 mM IPTG (EMAL283). We found that OptoLAC1B is able to keep YFP repressed under light conditions and effectively induce it when the culture is switched to the dark (Figure 5). Consistent with the time course experiments with the K-12 strain (Figure 2d), OptoLAC1B shows a ∼2 hour delay in protein expression (detectable by SDS-PAGE), whereas expression by IPTG induction is clearly observable within 1 hour of induction. Nevertheless, after 9-12 hours of induction in the dark, the levels of protein production achieved by OptoLAC1B are comparable to those achieved with IPTG induction. To determine the optimal cell density of induction, we examined protein expression at 9 hours after inducing at OD_600_ values ranging from 0.1-1.8 (Figure 5b). The maximum production of YFP is the same whether inducing with IPTG or darkness, and in both cases this maxima is achieved when inducing at low OD_600_ values (OD_600_ = 0.1-0.2). Furthermore, OptoLAC1B appears to maintain higher levels of YFP production when inducing at higher cell densities (OD_600_ > 1). These results demonstrate that our OptoLAC circuits can be robustly transferred across different strains and plasmids to adapt them to different biotechnological applications.

**Figure 5.**
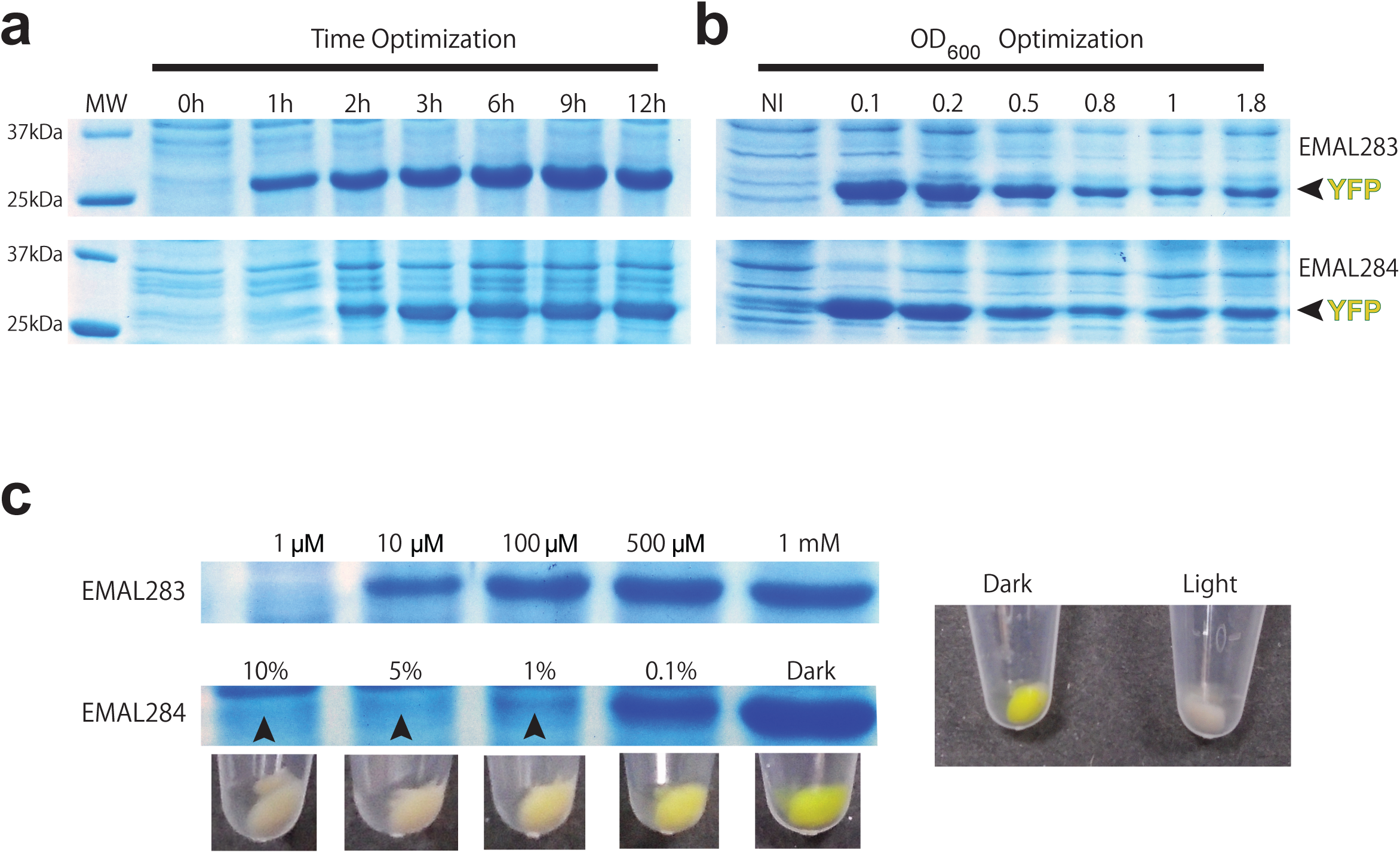
Light-controlled recombinant protein production in *E. coli* B strain (OptoBL). **(a)** Time course of recombinant YFP production. Protein production was induced at OD_600_ = 0.5 with either 1mM IPTG (EMAL283, top panel) or by switching cultures from light to dark using OptoLAC1B (EMAL284, bottom panel). **(b)** YFP production when inducing at different cell densities. Protein production was induced at OD_600_ = 0.1-1.8 with either 1mM IPTG (EMAL283, top panel), or by switching cultures from light to dark using OptoLAC1B (EMAL284, bottom panel). NI = Not induced: no IPTG added for EMAL283, kept under continuous blue light for EMAL284. **(c)** Tunability of YFP production using different doses of light or concentrations of IPTG, resolved with SDS-PAGE, or by visual inspection of bacterial pellets. Protein production was induced at OD_600_ = 0.5 with 1µM, 10µM, 100µM, 500µM, or 1mM IPTG (EMAL283, top panel), or by switching cultures to different doses of blue light using OptoLAC1B (EMAL284, bottom panel): 100 sec ON/900 sec OFF (10% light), 50 sec ON/950 sec OFF (5% light), 10 sec ON/990 sec OFF (1% light), 1 sec ON/999 sec OFF (0.1% light), and full darkness (0% light). All samples were resolved in SDS-PAGE (12% polyacrylamide gels).

Easier tunability of gene expression levels is an important advantage of optogenetic control over chemical induction. To showcase the tunability afforded by controlling light dose, we compared YFP production under different duty cycles of blue light (EMAL284) or different concentrations of IPTG (EMAL283). By varying IPTG concentrations from 1µM to 1mM, the increase in levels of YFP production shows a sigmoidal behavior, essentially saturating at IPTG concentrations above 100µM (Figure 5c). In contrast, varying light exposure from 0-10%, results in a more incremental response in the levels of YFP expression observed by both SDS-PAGE and visual inspection of the cell pellets (Figure 5c). Our findings illustrate the unique potential of light to precisely control the quantity and rate of protein production, both of which may impact the quality of recombinant proteins.

## Discussion

Our OptoLAC circuits offer unprecedented capabilities to control gene expression in *E. coli* with light. While several optogenetic tools have previously been developed for this organism^10,11,13^, our approach is the only one to harness the *lac* operon, which allows for the direct substitution of IPTG with light as an inducing agent for gene expression. Our results show the feasibility of such replacement for at least two common applications: metabolic engineering and recombinant protein production (Table 1). All the existing IPTG-inducible vectors that we tested, which use five different *lacO*-containing promoters (P_T5-lacO_, P_lacUV5_, P_trc_, P_L_lacO1_, and P_T7_), were easily adapted to work with our OptoLAC circuits and *lacI*-strains (OptoMG or OptoBL) for optogenetic control. The only requirements are that any copy of *lacI* is removed from the existing vectors, and that they use a unique selection marker and origin of replication. OptoLAC1B (pMAL658), which contains a different origin and selection marker than our other OptoLAC circuits (pMAL294, pMAL296, and pMAL630), offers some flexibility in these last two requirements.

**Table 1:**
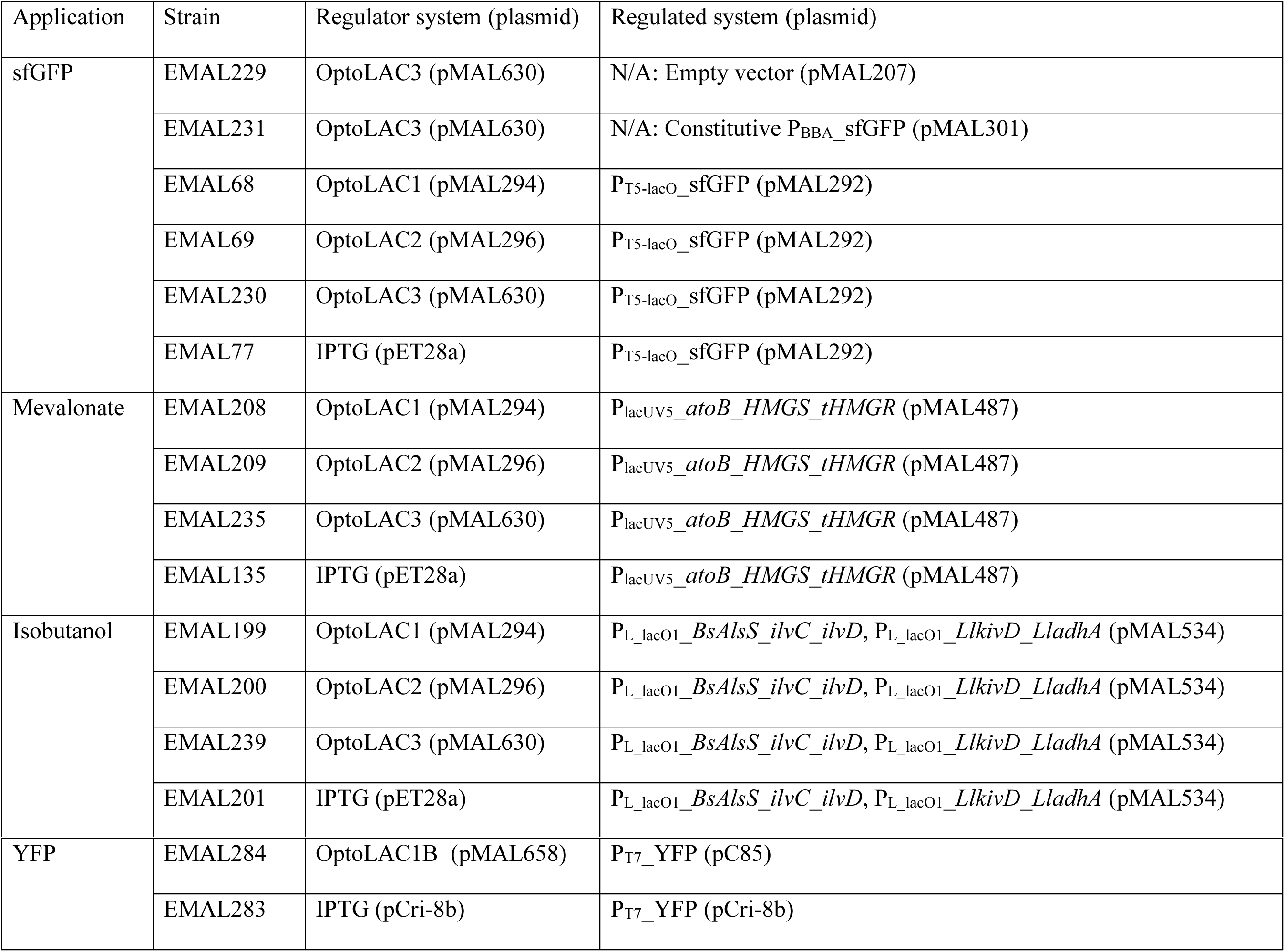
Strain breakdown for each application

| Application | Strain | Regulator system (plasmid) | Regulated system (plasmid) |
| --- | --- | --- | --- |
| sfGFP | EMAL229 | OptoLAC3 (pMAL630) | N/A: Empty vector (pMAL207) |
|  | EMAL231 | OptoLAC3 (pMAL630) | N/A: Constitutive P<sub>BBA</sub>_sfGFP (pMAL301) |
|  | EMAL68 | OptoLAC1 (pMAL294) | P<sub>T5-lacO</sub>_sfGFP (pMAL292) |
|  | EMAL69 | OptoLAC2 (pMAL296) | P<sub>T5-lacO</sub>_sfGFP (pMAL292) |
|  | EMAL230 | OptoLAC3 (pMAL630) | P<sub>T5-lacO</sub>_sfGFP (pMAL292) |
|  | EMAL77 | IPTG (pET28a) | P<sub>T5-lacO</sub>_sfGFP (pMAL292) |
| Mevalonate | EMAL208 | OptoLAC1 (pMAL294) | P<sub>lacUV5_atoB_HMGS_tHMGR</sub> (pMAL487) |
|  | EMAL209 | OptoLAC2 (pMAL296) | P<sub>lacUV5_atoB_HMGS_tHMGR</sub> (pMAL487) |
|  | EMAL235 | OptoLAC3 (pMAL630) | P<sub>lacUV5_atoB_HMGS_tHMGR</sub> (pMAL487) |
|  | EMAL135 | IPTG (pET28a) | P<sub>lacUV5_atoB_HMGS_tHMGR</sub> (pMAL487) |
| Isobutanol | EMAL199 | OptoLAC1 (pMAL294) | P<sub>L_lacO1_BsAlsS_ilvC_ilvD</sub>, P<sub>L_lacO1_LlkivD_LladhA</sub> (pMAL534) |
|  | EMAL200 | OptoLAC2 (pMAL296) | P<sub>L_lacO1_BsAlsS_ilvC_ilvD</sub>, P<sub>L_lacO1_LlkivD_LladhA</sub> (pMAL534) |
|  | EMAL239 | OptoLAC3 (pMAL630) | P<sub>L_lacO1_BsAlsS_ilvC_ilvD</sub>, P<sub>L_lacO1_LlkivD_LladhA</sub> (pMAL534) |
|  | EMAL201 | IPTG (pET28a) | P<sub>L_lacO1_BsAlsS_ilvC_ilvD</sub>, P<sub>L_lacO1_LlkivD_LladhA</sub> (pMAL534) |
| YFP | EMAL284 | OptoLAC1B (pMAL658) | P<sub>T7</sub>_YFP (pC85) |
|  | EMAL283 | IPTG (pCri-8b) | P<sub>T7</sub>_YFP (pCri-8b) |

By design, our OptoLAC circuits are inducible by darkness to alleviate potential challenge of light penetration at the high cell densities that often occur during the production phase of fermentation. This feature might have played a role in our successful scale-up experiments in a 2-L bioreactor. However, it is important to note that light penetration might not be as limiting to our systems as it is, for example, to cultures of photosynthetic organisms (e.g. algae or cyanobacteria), where there is a stoichiometric relationship between the photons absorbed by the photosynthetic systems and ATP output^32–35^. The light harvesting antenna in photosynthetic organisms present an additional challenge to effectively illuminate cultures for robust cell growth and production. In contrast, our optogenetic circuits are not required for energy metabolism or expressed in an organism evolved to compete for light harvesting. Instead, they drive a genetic program to control the expression of the cI repressor, leading to the relatively long-lived accumulation of LacI mRNA and protein under blue light, which does not immediately subside when the cells are switched to the dark. In this sense, our optogenetic circuits have an inherent *memory* of having been exposed to blue light, which makes the requirements for light penetration more lenient.

This inherent *memory* in our system is evident in the kinetics experiments (Figure 2c), bioreactor fermentations (Figure 4), and protein production (Figure 5). In all these instances, the response of our circuits to changing conditions from light to darkness has a ∼2 hour delay, as opposed to the almost immediate response to IPTG (Figure 2c) caused by its mechanism of action as a LacI protein inhibitor. This feature of our optogenetic circuits not only alleviates potential challenges to light penetration, but might also be useful to mimic slow-acting inducible systems, such as those based on quorum sensing, which have been effectively used as auto-inducible systems for chemical production^36–38^. Gradual decrease of illumination could be used to mirror the gradual increase of quorum-sensing molecule concentrations, but with the advantage that the onset and rate of light dimming can be easily controlled and operated reversibly using our OptoLAC circuits.

Despite the time delay in our optogenetic circuits, they eventually reach or exceed the expression levels achieved with IPTG induction. By the sixth hour after induction, OptoLAC1 and OptoLAC2 surpass the level of GFP expression reached with IPTG (Figure 2d). Furthermore, mevalonate and isobutanol fermentations achieve higher titers when controlled with our optogenetic circuits than with IPTG induction (using the same plasmid containing the metabolic pathways). This suggests that, at least in some conditions, the cI-mediated repression of *lacI* achieved by our circuits is more effective at allowing gene expression from *lacO*-containing promoters than IPTG is at sustaining full inhibition of LacI.

Our initial suite of OptoLAC circuits adds flexibility to optogenetic control of engineered metabolic pathways. When producing mevalonate, OptoLAC1 and OptoLAC2 achieve approximately the same maximum titers (6.4 ± 0.2 and 6.1 ± 0.2 g/L, respectively) with approximately the same relatively low optimal ρ_s_ values (0.17 and 0.23, respectively). Given the low toxicity of mevalonate, it is not surprising that its production would benefit from inducing early in the fermentation at low OD_600_. In contrast, for isobutanol production, OptoLAC1 and OptoLAC3 also achieve equal maximum titers (2.5 ± 0.1 and 2.4 ± 0.2 g/L, respectively), but at substantially different ρ_s_ values (1.00 and 0.24, respectively), which reflects the ability of our circuits to adapt to the relatively higher toxicity of isobutanol. In the case of OptoLAC1, which is stronger but also leakier than OptoLAC3, the maximum titers are achieved when inducing at higher OD_600_, probably because of the accumulation of toxic product during the growth phase. On the other hand, OptoLAC3 is not as strong as OptoLAC1 but is better at keeping isobutanol biosynthesis tightly repressed in the light, which may explain why the maximum isobutanol titers are achieved when the fermentation is induced earlier, at lower OD_600_. These observations highlight the benefit of having a flexible suite of OptoLAC circuits with different strengths, sensitivities, and folds of induction, which could make each of them more suitable for different metabolic pathways or biotechnological applications.

The fine-tunability of optogenetic systems may be advantageous to optimally balance metabolic pathways for chemical production. The levels and ratios at which different metabolic enzymes in an engineered pathway are expressed can have profound effects on the availability of enzyme substrates and cofactors, the accumulation of byproducts or toxic intermediates, and the metabolic burden on the cell, thus impacting the yield and productivity of the pathway^39^. This key challenge in metabolic engineering is currently addressed by assembling a large number of constructs, usually in biofoundries^40^, to combinatorially vary the levels of expression of different enzymes in the pathway of interest^41–43^. Our results show that OptoLAC circuits can tune and control the levels of gene expression of different promoters and in different media in fermentations at least as long as 72 hours (Figure 2a-c), which could greatly reduce the number of constructs needed to find the optimal balance of enzyme expression in a metabolic pathway. Furthermore, our blue light-responsive OptoLAC system could be combined with future circuits applied to metabolic engineering that respond orthogonally to different wavelengths of light (e.g. green, red, or infrared^12,14,44,45^). This could allow the application of optogenetics to the powerful multivariate-modular approach to engineer balanced metabolic pathways^46^, by tuning two or more modules of a pathway simultaneously with different colors of light. Such a polychromatic approach would not only dramatically reduce the number of constructs required to balance complex metabolic pathways, relative to current practices, but also increase the resolution at which levels of enzyme expression could be screened for pathway optimization.

In addition to showing the efficacy of our optogenetic *lac* operon for metabolic engineering, we also demonstrate its applicability in recombinant protein production. This confirms the ability to use OptoLAC circuits not only in K-12 strains, for chemical production, but also in B strains for protein expression. Light control of gene expression may become advantageous when complications in recombinant protein production arise. For example, if the protein being produced is toxic to *E. coli*, it often leads to low yields or makes it necessary to add inhibitors of the toxic protein during production^47^. Another common complication occurs when the rates of protein expression obtained by IPTG-induction, even at low IPTG concentrations, are too high to keep the protein from aggregating or forming inclusion bodies. This problem can sometimes be alleviated by inducing with IPTG at low temperatures^48^ or by inducing slowly using autoinduction media^49^, but more often it altogether prevents the expression of functional recombinant protein. It is possible that the fine-tunability and reversibility of gene expression afforded by light could enhance the yields of toxic proteins by tempering their levels of expression, or improve the quality and solubility of proteins that are difficult to produce by regulating their rates of expression.

Our OptoLAC circuits open the door to using light control as a replacement for IPTG induction for a broad number of applications. In industrial fermentations, eliminating the use IPTG (or other chemical inducers) would reduce costs and increase production. Additionally, it would enable dynamic controls in processes for which the use of chemical inducers is prohibitively expensive. Beyond chemical and recombinant protein production, IPTG induction has been used for a variety of research applications, including to study persistence^50^, bacterial motility and swarming^51^, inducible protein degradation^52^, biofilm formation^53^, and biomaterials^54^. Furthermore, the *lac* operon has been exported to other bacteria for IPTG inducibility^55–60^. Therefore, our OptoLAC circuits may be applicable in a wide range of systems originally designed to be IPTG-inducible, providing them with the enhanced capabilities typically offered by light, such as fine-tunability, reversibility, and spatiotemporal control.

## Online Methods

### Plasmid construction

Plasmids were cloned into *E. coli* strain DH5α made chemically competent using the Inoue method^61^. Transformants were inoculated on LB agar plates with appropriate antibiotics: 100 µg/mL ampicillin, 100 µg/mL carbenicillin, 50 µg/mL kanamycin, 34 µg/mL chloramphenicol, or 50 µg/mL spectinomycin. Epoch Life Science Miniprep, Omega Gel Extraction, and Omega PCR purification kits were used to extract and purify plasmids and DNA fragments. Backbones and inserts were either digested using restriction enzymes purchased from NEB or PCR amplified using Q5 Polymerase from NEB or CloneAmp HiFi PCR premix from Takara Bio. One-step Gibson isothermal assembly reactions were performed based on previously described protocols^62^. Primers were ordered from Integrated DNA Technologies (Coralville, IA) or Genewiz (South Plainfield, NJ). All plasmids were verified using Sanger sequencing from Genewiz. We avoid using tandem repeats to prevent recombination after transformation, and thus do not observe instability of strains or plasmids.

Promoters, operators, and tags (P_BBa_J23100_, P_lacUV5_, P_trc_, *lacO*, ssRA tags) were inserted by Gibson assembly, using extended-length primers with homology arms. Plasmids pDusk and pDawn were obtained as a gift from Andreas Möglich (Addgene #43795 and 43796, respectively)^15^. Plasmid pMevT, used for mevalonate production, was obtained as a gift from Jay Keasling (Addgene #17815)^24^. Plasmids pSA65 and pSA69, used for isobutanol production, were obtained as gifts from James Liao^27,63^. A detailed description of all the plasmids used in this study can be found in Supplementary Table 1.

### Bacterial cell culture growth and measurements

Unless otherwise specified, liquid *E. coli* cultures were inoculated from single colonies on LB + antibiotic agar plates and grown in 96-well (USA Scientific Item #CC7672-7596) or 24-well (USA Scientific Item #CC7672-7524) plates in a New Brunswick Innova 4000 incubator shaker set to 37°C and shaken at 200 RPM (19 mm orbital diameter). We used M9 minimal salts media supplemented with 0.2% w/v casamino acids (Bio Basic), K3 trace metal mixture^64^, 2% w/v (20 g/L) glucose, and appropriate antibiotics (using the previously specified concentrations), unless stated otherwise. To stimulate cells with blue (465 nm) light, we used LED panels (HQRP New Square 12” Grow Light Blue LED 14W) placed above the culture such that light intensity was between 80-110 µmol/m^2^/s as measured using a Quantum meter (Apogee Instruments, Model MQ-510). To control light duty cycles, LED panels were regulated with a Nearpow Multifunctional Infinite Loop Programmable Plug-in Digital Timer Switch.

To measure cell concentration, optical density measurements were taken at 600 nm (OD_600_), using media (exposed to the same light and incubation conditions as the bacteria cultures) as blank. Measurements were taken using a TECAN plate reader (infinite M200PRO) or Eppendorf spectrophotometer (BioSpectrometer basic) with a microvolume measuring cell (Eppendorf µCuvette G1.0), using samples diluted to a range of OD_600_ between 0.1 and 1.0.

### Flow Cytometry

Super-folder GFP (GFP) expression was quantified by flow cytometry using a BD LSR II flow cytometer (BD Biosciences, San Jose, CA, USA) with an excitation wavelength of 488 nm and an emission wavelength of 530 nm. The gating used in our analyses was defined to include positive (EMAL231) and negative (EMAL229) control cells based on GFP fluorescence, but exclude particles that are either too small or too large to be single living bacterial cells, based on the side scatter (SSC-A) vs forward scatter (FSC-A) plots as previously described^7^. Median fluorescence values were determined from 10,000 cells.

The fluorescence data was normalized against the background fluorescence from cells lacking GFP (EMAL229) to account for potential light bleaching and cell auto-fluorescence. All fluorescence measurements were either taken at the end of the experiments or on aliquots taken from experimental cultures so that potential activation of YF1-FixJ by the light used to excite GFP did not affect our experiments or results.

### Fermentation sample preparation and analytical methods

For mevalonate production, 560 µL of cell culture was mixed with 140 µL of 0.5 M HCl in a 1.5 mL microcentrifuge tube and vortexed at high speed for 1 minute to convert mevalonate to (±)-mevalonolactone. The mixture was then centrifuged at 17,000 RCF for 45 minutes at 4°C in a benchtop centrifuge (Eppendorf Centrifuge 5424) to obtain cell-free supernatant. Then, 250 µL of this supernatant was transferred to an HPLC vial for analysis. For isobutanol production, 700 µL of cell culture was transferred to a 2 mL microcentrifuge tube and centrifuged as described above to obtain cell-free supernatant. Then, 250 µL of this supernatant was transferred to an HPLC vial for analysis.

Cell-free supernatant samples were analyzed via liquid chromatography (Agilent 1260 Infinity) using an Aminex HPX-87H ion-exchange column (Bio-Rad, Richmond, CA, USA). The mobile phase was 5 mM sulfuric acid. Samples were run through the column at 55°C and a flow rate of 0.6 ml/min. Glucose, mevalonate, and isobutanol were monitored with a refractive index detector (RID, Agilent G1362A). To determine their concentration, the peak areas were measured and compared to those of their own standard solutions for quantification, using (±)-mevalonolactone (Sigma-Aldrich) for mevalonate.

### Construction of OptoLAC circuits

Deletion of *lacI* was performed using the Datsenko-Wanner method^65^, using primers with 70 base pairs of homology to the promoter and terminator regions of the targeted gene. These primers were used to amplify the kanamycin resistance marker flanked by FRT sites from pKD4, or the spectinomycin resistance marker from pYTK91^66^. Cells were made electro-competent and electroporated as described previously^67^. FRT-flanked resistance markers were cured using FLP recombinase from pCP20^68^. Gene deletions were genotyped by sequencing PCR products amplified from purified genomic DNA using primers flanking the region of deletion. A detailed description of all the strains used in this study can be found in Supplementary Table 2.

Strain MG1655 *ΔlacI::FRT-KanR-FRT* was obtained as a gift from Mark Brynildsen^17^. From this strain, we removed the kanamycin resistance marker remaining from *lacI* deletion through FLP-FRT recombination. We hereafter call this strain OptoMG (EMAL52) in this study. Chemically competent OptoMG was transformed with pDawn controlling *lacI* (pMAL288) and a reporter plasmid containing P_T5-lacO_-GFP (pMAL292) to make EMAL57. Three different ssRA tags with progressively weaker degradation strengths^21^ (based on the last three residues) were added to the C terminus of *lacI*: LAA (pMAL294/OptoLAC1), AAV (pMAL296/OptoLAC2), and ASV (pMAL300). OptoMG was transformed with pMAL292 and each of these plasmids to make EMAL68, EMAL69, and EMAL71, respectively. To construct OptoLAC3, a *lacO* site was added between the P_FixK2_ promoter and its downstream ribosome-binding site in pMAL296 (Supplementary Sequence 2), creating pMAL630. OptoMG was transformed with pMAL292 and pMAL630 to make EMAL230.

As controls for GFP measurement, we transformed OptoMG with pMAL630 (OptoLAC3) and pMAL207 (Empty vector), to produce EMAL229. This strain serves as a negative control for subtraction of cell auto-fluorescence. In addition, we transformed OptoMG with pMAL630 (OptoLAC3) and pMAL301 (P_BBA_-GFP), to produce EMAL231. This strain expresses GFP constitutively, serving as a positive control to check for photobleaching of GFP under blue light.

### Characterization of OptoLAC circuits

To evaluate the response of our circuits to blue light, we grew 1 mL overnight cultures of EMAL57, EMAL68, EMAL69, EMAL71, EMAL229, EMAL230, and EMAL231. These overnight cultures were grown under blue light to avoid premature transcription of GFP. We back-diluted the cultures to OD_600_ = 0.01 in 150 µL triplicates into separate 96-well plates and grew the cultures for 8 hours under blue light or in the dark (by wrapping the plate in aluminum foil). After growing for 8 hours, 1 µL from each well was diluted into separate wells containing 199 µL of ice-cold phosphate buffered saline (Corning Life Sciences) and taken for flow cytometry analysis. For light dose-response analysis, additional conditions of light were tested by applying light pulses of 10s ON/990s OFF and 100s ON/900s OFF.

To test the performance of our circuits with different IPTG-inducible promoters, chemically competent OptoMG was transformed with pMAL294 (OptoLAC1) and one of the following two plasmids: a minimal-copy reporter plasmid containing P_lacUV5_-GFP (pMAL302) or P_trc_-GFP (pMAL303). We named these strains EMAL267 and EMAL268, respectively. Overnight cultures of 1 mL of EMAL229, EMAL231, EMAL267, and EMAL268 (grown under blue light) were back-diluted to OD_600_ = 0.01 in 150 µL triplicates into separate 96-well plates and grown for 8 hours under blue light, pulses of 10s ON/990s OFF and 100s ON/900s OFF, or in the dark (by wrapping the plate in aluminum foil). After growing for 8 hours, 1 µL from each well was diluted into separate wells containing 199 µL of ice-cold phosphate buffered saline and taken for flow cytometry analysis.

To test different carbon sources, 1 mL overnight cultures of EMAL229, EMAL231, and EMAL68 (grown under blue light) were grown in M9 + 20 g/L glucose or xylose, LB (Miller) or SOB media. We back-diluted the cultures to OD_600_ = 0.01 in 150 µL of the same overnight media in triplicates into separate 96-well plates, and grew the cultures for 8 hours under full blue light, light pulses of 10s ON/990s OFF or 100s ON/900s OFF, or in the dark (by wrapping the plate in aluminum foil). After growing for 8 hours, 1 µL from each well was diluted into separate wells containing 199 µL of ice-cold phosphate buffered saline and taken for flow cytometry analysis.

To test spatial control of GFP expression, we grew 1 mL overnight cultures of EMAL68 under blue light. 500 µL of culture was spread onto a 150mm x 15mm LB + kanamycin + ampicillin agar plate (Laboratory Disposable Products, Inc. Catalog #229656-CT) using sterile glass beads. The plate was placed in ambient temperature on top of a black cloth and 35cm underneath a projector (Epson H764A) displaying an image of a tiger (Drawing Work). The setup was covered in black cloth to prevent ambient light contamination. After 16 hours, the plate was imaged for GFP fluorescence using a Bio-Rad ChemiDoc MP Imaging System with Image Lab software, with Blue Epi illumination using a 530/28 Filter (Filter 4).

### Kinetic analysis of OptoLAC circuits

We prepared an IPTG-inducible control strain that contains constitutively expressed *lacI* by transforming OptoMG with pMAL292 and pET28a, creating strain EMAL77. We inoculated 1 mL overnight cultures of EMAL229, EMAL68, EMAL69, EMAL230, and EMAL77 (see Table 1). Cells were grown under blue light as described above to avoid premature transcription of GFP. We back-diluted the cultures to OD_600_ = 0.01 in 1 mL triplicates into 8 separate 24-well plates and grew the cultures as described above for approximately 4 hours under blue light, at which point the cultures reached OD_600_ = 0.9. At this point, we collected samples from one plate (corresponding to t = 0 hours). For strains EMAL229, EMAL68, EMAL69, and EMAL230, we induced 6 of the remaining 7 plates by wrapping them in aluminum foil. For EMAL77, we instead induced by adding IPTG to 6 of the remaining 7 plates, to a final concentration of 1mM. The final (8^th^) plate was left in blue light (for EMAL77, without IPTG) for 12 hours as a control. Every 2 hours up to 12 hours after induction one plate from each strain was uncovered to measure the levels of GFP expression (and then discarded), by diluting 1 µL of culture into separate wells containing 199 µL of ice-cold phosphate buffered saline (Corning Life Sciences) for flow cytometry analysis.

### Construction of mevalonate producing strains

We transformed chemically competent OptoMG cells with either OptoLAC1, OptoLAC2 or OptoLAC3 and a modified version of pMevT^24^ in which P_lac_ was replaced with P_lacUV5_ (pMAL487), generating strains EMAL208, EMAL209, and EMAL235, respectively. A light-insensitive IPTG-inducible control strain was made by transforming chemically competent OptoMG cells with pMAL487 and pET28a, generating EMAL135 (see Table 1). These strains (EMAL208, EMAL209, EMAL235, and EMAL135) express, from a single operon, the first three enzymes of the mevalonate pathway, which convert acetyl-CoA into mevalonate: acetoacetyl-CoA thiolase (*ato*B) from *E. coli*, 3-hydroxy-3-methylglutaryl-CoA synthase (HMGS or *ERG13*) from *Saccharomyces cerevisiae*, and a truncated version of 3-hydroxy-3-methylglutaryl-CoA reductase (tHMGR) from *S. cerevisiae*. The transformants were plated on LB + kanamycin + chloramphenicol agar and colonies were grown under blue light to avoid potential negative selection due to pathway expression. Four colonies from each strain were screened for mevalonate production. Each colony was used to inoculate 1 mL of M9 + 2% glucose + chloramphenicol + kanamycin media, grown in a 24-well plate overnight at 37°C and 200 RPM under blue light. The next day, each culture was back-diluted to an OD_600_ of 0.01 and grown for 4 hours at 37°C at 200 RPM under blue light. The plates were then sealed with Nunc Sealing Tape (Thermo Scientific) and wrapped in aluminum foil; for the control strain EMAL135, IPTG was added to 1 mM before sealing and wrapping. A single hole (using a sterile syringe needle) was poked in the center of the sealing tape covering each sample well to allow limited aeration. The strains were fermented in the dark at 37°C at 200 RPM for 72 hours, after which samples were prepared for HPLC analysis as described above. The highest producing colonies were selected for subsequent optimization.

To find the optimal value of ρ_s_ for mevalonate production, we back-diluted 1 mL overnight cultures of each strain to different OD_600_ values (ranging from 0.001 to 0.1) in 1 mL cultures of the same media described above (in quadruplicates). The different dilutions were grown for 4 hours, reaching different OD_600_ values, at which time cultures were switched from light to dark conditions (values in Figure 3b,c) or induced with IPTG (Supplementary Figure 3a). As controls, we grew each strain under continuous blue light (EMAL208, EMAL209, EMAL235) or with no IPTG added (EMAL135). The same fermentation procedure described above was then followed.

### Construction of isobutanol producing strains

We transformed chemically competent OptoMG cells with either OptoLAC1, OptoLAC2 or OptoLAC3 and a plasmid that combines all the genes from pSA65 and pSA69 in a single vector with a p15A origin of replication and β-lactamase resistance marker (pMAL534), generating strains EMAL199, EMAL200, and EMAL239, respectively. A light-insensitive IPTG-inducible control strain was made by co-transforming chemically competent OptoMG with pMAL534 and pET28a, generating EMAL201 (see Table 1). These strains (EMAL199, EMAL200, EMAL239, and EMAL201) express from the P_L_lacO1_ promoter three enzymes which convert pyruvate to 2-ketoisovalerate: acetolactate synthase from *Bacillus subtilis* (Bs*AlsS*), acetohydroxyacid reductoisomerase (*ilvC*), and dihydroxyacid dehydratase (*ilvD*). They also express from another P_L_lacO1_ promoter two enzymes that convert 2-ketoisovalerate into isobutanol: 2-ketoacid decarboxylase from *Lactococcus lactis* (*Llkivd)* and alcohol dehydrogenase from *Lactococcus lactis* (*LladhA*). The transformants were plated on LB + kanamycin + ampicillin agar and colonies were grown under blue light to avoid negative selection due to the potential pathway expression. Four colonies from each strain were screened for isobutanol production. Each colony was used to inoculate 1 mL of M9 + 2% glucose + kanamycin + carbenicillin media, grown in a 24-well plate overnight at 37°C and 200 RPM under blue light. The next day, each culture was back-diluted in the same media to an OD_600_ of 0.01 and grown for 4 hours at 37°C at 200 RPM under blue light. The plates were then sealed with Nunc Sealing Tape (Thermo Scientific) and wrapped in aluminum foil; for the control strain EMAL201, IPTG was added to 1 mM before sealing and wrapping. No holes were poked in the sealing tape to prevent evaporation of isobutanol. The strains were fermented in the dark at 30°C at 200 RPM for 72 hours, after which samples were prepared for HPLC analysis as described above. The highest producing colonies were selected for subsequent optimization.

To find the optimal value of ρ_s_ for isobutanol production, we back-diluted 1 mL overnight cultures of each strain to different OD_600_ values (ranging from 0.001 to 0.1) in 1 mL cultures of the same media described above (in quadruplicates). The different dilutions were grown for 4 hours, reaching different OD_600_ values, at which time cultures were switched from light to dark conditions (values in Figure 3e,f), or induced with IPTG (Supplementary Figure 3b). As controls, we grew each strain under continuous blue light (EMAL199, EMAL200, EMAL239) or with no IPTG added (EMAL201). The same fermentation procedure described above was then followed.

### Scale-up of chemical production in a 2-L bioreactor

To validate our circuits for chemical production in higher cell density conditions and larger culture volumes, we grew a 5 mL overnight culture of EMAL208 (OptoLAC1 driving mevalonate production) in M9 + 5% glucose + kanamycin + chloramphenicol under blue light at 37°C. We then set up a BioFlo120 system with a 2-L bioreactor (Eppendorf, B120110001) and added 1 L of autoclaved M9 + 5% glucose + kanamycin + chloramphenicol . The reactor was set to 37°C, pH of 7.0 (maintained using a base feed of 2 M potassium hydroxide), and a minimum dissolved oxygen percentage of 20 (maintained by adjusting the agitation rate between 200-800 RPM and by injecting air at a flow rate of 0.1-3.0 SLPM). Three blue LED panels were placed in triangular formation ∼20 cm from the reactor such that the light illuminated approximately 61.5% of the bulk surface area at an intensity of 80-110 µmoles/m^2^/s. The reactor was then inoculated to an OD_600_ of 0.004, and the cells were grown for about 4 hours under blue light until reaching an OD_600_ of 0.17. The lights were then turned off and the reactor was wrapped in aluminum foil and covered with black cloth. At 8 hours, 50 µL of Antifoam 204 (Sigma-Aldrich) was added to prevent foaming. Culture samples of 1 mL were taken while minimizing potential exposure of the culture to ambient light and prepared for HPLC analysis as described above.

### Protein production

To develop strains for protein production, we knocked out the endogenous copy of *lacI* from BL21 DE3 and removed the remaining kanamycin resistance marker left by the knockout (as described above), resulting in EMAL224. We then deleted the copy of *lacI* introduced by the DE3 prophage, resulting in EMAL255. We hereafter call this strain OptoBL in this study. We transformed electrocompetent OptoBL cells with pMAL658 (OptoLAC1B) and pC85 (P_T7_-YFP, P_lacIQ__*lacI*), resulting in EMAL284. As an IPTG-inducible control containing constitutive *lacI*, we transformed BL21 DE3 with pCri-8b^30^, resulting in EMAL283.

For kinetic analysis of protein production, we inoculated 1 mL overnight cultures of EMAL283 and EMAL284 into LB Miller + kanamycin (+ chloramphenicol for EMAL284). EMAL284 cells were grown under blue light to avoid premature transcription of YFP. We back-diluted the cultures to OD_600_ = 0.05 in 1 mL cultures into eight separate 24-well plates and grew the cultures to an OD_600_ = 0.5 (which took approximately 3 hours) under blue light. We then switched 7 of the plates to the dark by wrapping them in aluminum foil, adding IPTG to a final concentration of 1mM for EMAL283. One plate was left in blue light and without IPTG for 12 hours as a control. The OD_600_ of each sample was measured before harvesting the full 1 mL culture at 0 hours, 1 hour, 2 hours, 3 hours, 6 hours, 9 hours, and 12 hours. Samples were harvested by centrifuging at 17,000 RCF for 5 minutes in a benchtop microcentrifuge (Thermo Scientific, Sorvall Legend Micro 17) at room temperature. The pellets were then resuspended in 200 µL of resuspension buffer (Tris 50 mM, pH = 8.0 and 300 mM NaCl), mixed with 50 µL of SDS sample buffer, and incubated at 100°C in a heat block (Eppendorf ThermoMixer C) for 10 min at 700 rpm. Samples were loaded onto 12% SDS-PAGE gels and resolved. The volumes loaded on the gels (between 5 to 25 µL) were adjusted based on the OD_600_ measured just before the samples were harvested, in order to load approximately the same amount of total protein for each corresponding time point. Gels were stained with Coomassie Brilliant Blue G-250.

To find the optimal ρ_s_ value for YFP production, we inoculated 1 mL overnight cultures of EMAL283 and EMAL284 into LB Miller + kanamycin (+ chloramphenicol for EMAL284). EMAL284 cells were grown under blue light to avoid premature transcription of YFP. We back-diluted the cultures to OD_600_ values between 0.001 and 0.1 in 1 mL cultures into a 24-well plate and grew the cultures to OD_600_ values between 0.1 and 1.8 (which took approximately 3 hours) under blue light. We then induced YFP expression by switching the plate to the dark (wrapping it in aluminum foil), or adding IPTG to a final concentration of 1mM for EMAL283. The OD_600_ of each sample was measured before harvesting the full 1 mL culture at 9 hours as previously described.

To test tunability of YFP production using different exposures to light or IPTG concentrations, we inoculated 1 mL overnight cultures of EMAL283 and EMAL284 into LB Miller + kanamycin (+ chloramphenicol for EMAL284). EMAL284 cells were grown under blue light to avoid premature transcription of YFP. For EMAL284, we back-diluted the cultures to OD_600_ = 0.05 in 1 mL cultures into five separate 24-well plates and grew the cultures to an OD_600_ = 0.5 (which took approximately 3 hours) under blue light. We then moved the plates to five separate light conditions: pulses of 1s ON/999s OFF, 10s ON/990s OFF, 50s ON/950s OFF, 100s ON/900s OFF, or in the dark (by wrapping the plate in aluminum foil). For EMAL283, we back-diluted the cultures to OD_600_ = 0.05 in 1 mL quintuplicates into a 24-well plate and grew the cultures to an OD_600_ = 0.5 (which took approximately 3 hours). We then added IPTG to final concentrations of 1µM, 10µM, 100µM, 500µM, and 1mM. The OD_600_ of each sample was measured before harvesting the full 1 mL culture at 9 hours as previously described.

### Statistics

Statistical significance was determined using a standard t-test for P values. T scores were calculated by the formula: 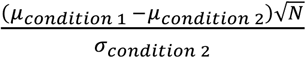. P values were calculated using a degree of freedom of 2 and a two-sided t-test calculator.

### Data Availability

The authors declare that all data supporting the findings of this study are available within the paper (and its supplementary information files), but original data that supports the findings are available from the corresponding authors upon reasonable request.

## Acknowledgements

We are immensely grateful to Wendy Mok and Mark Brynildsen for advice and troubleshooting with *E. coli* protocols. We thank Dr. Christina DeCoste, Dr. Katherine Rittenbach, and the Princeton Molecular Biology Flow Cytometry Resource Center for assistance with flow cytometry experiments. J.L.A was supported by the U.S. Department of Energy, Office of Science, Office of Biological and Environmental Research Award Number DE-SC0019363, the NSF CAREER Award CBET-1751840, The Pew Charitable Trusts, The Eric and Wendy Schmidt Transformative Technology Fund Award, and the Camille Dreyfus Teacher-Scholar Award.

## Author contributions

M.A.L. and J.L.A. conceived this project and designed the experiments. M.A.L. and S.S.I. constructed the strains and plasmids. M.A.L., S.S.I., C.C-L., E.M.Z., and H.K. executed the experiments. M.A.L. and J.L.A. analyzed the data and wrote the paper.

## Supplementary Information

### Supplementary Sequences

#### Supplementary Sequence 1

##### P_FixK2_

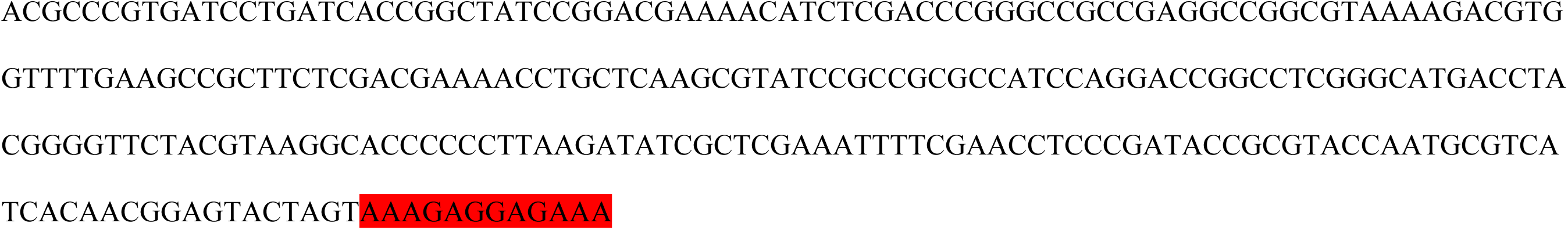

#### Supplementary Sequence 2

### P_FixK2_lacO_

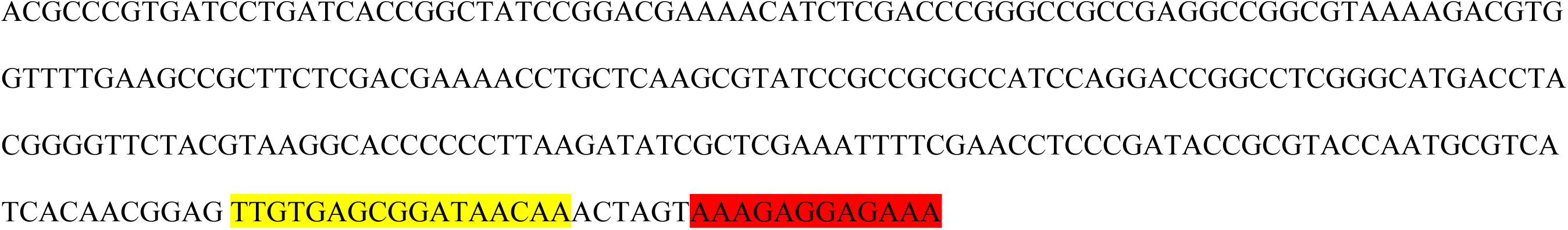

**Supplementary Table 1:**
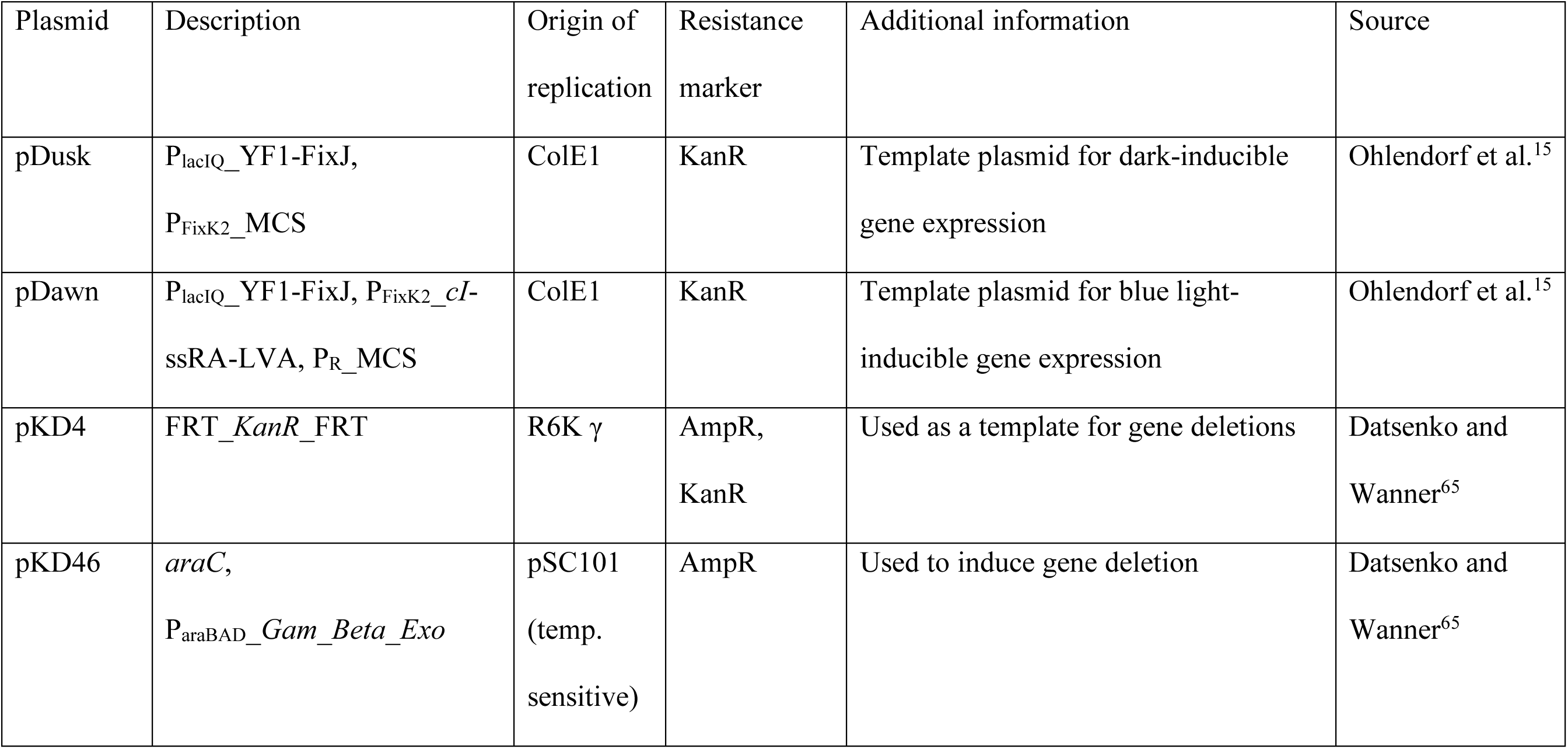
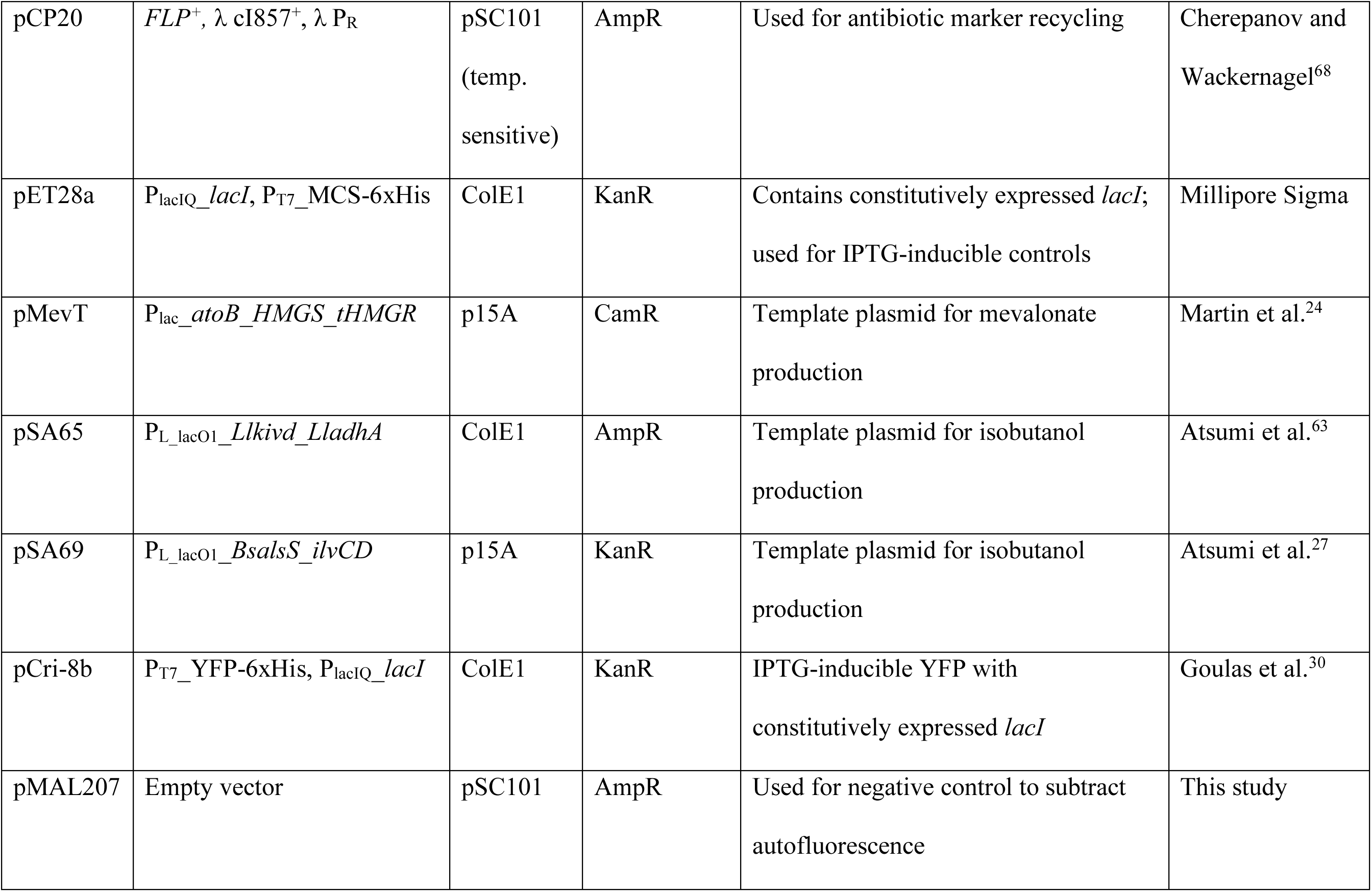
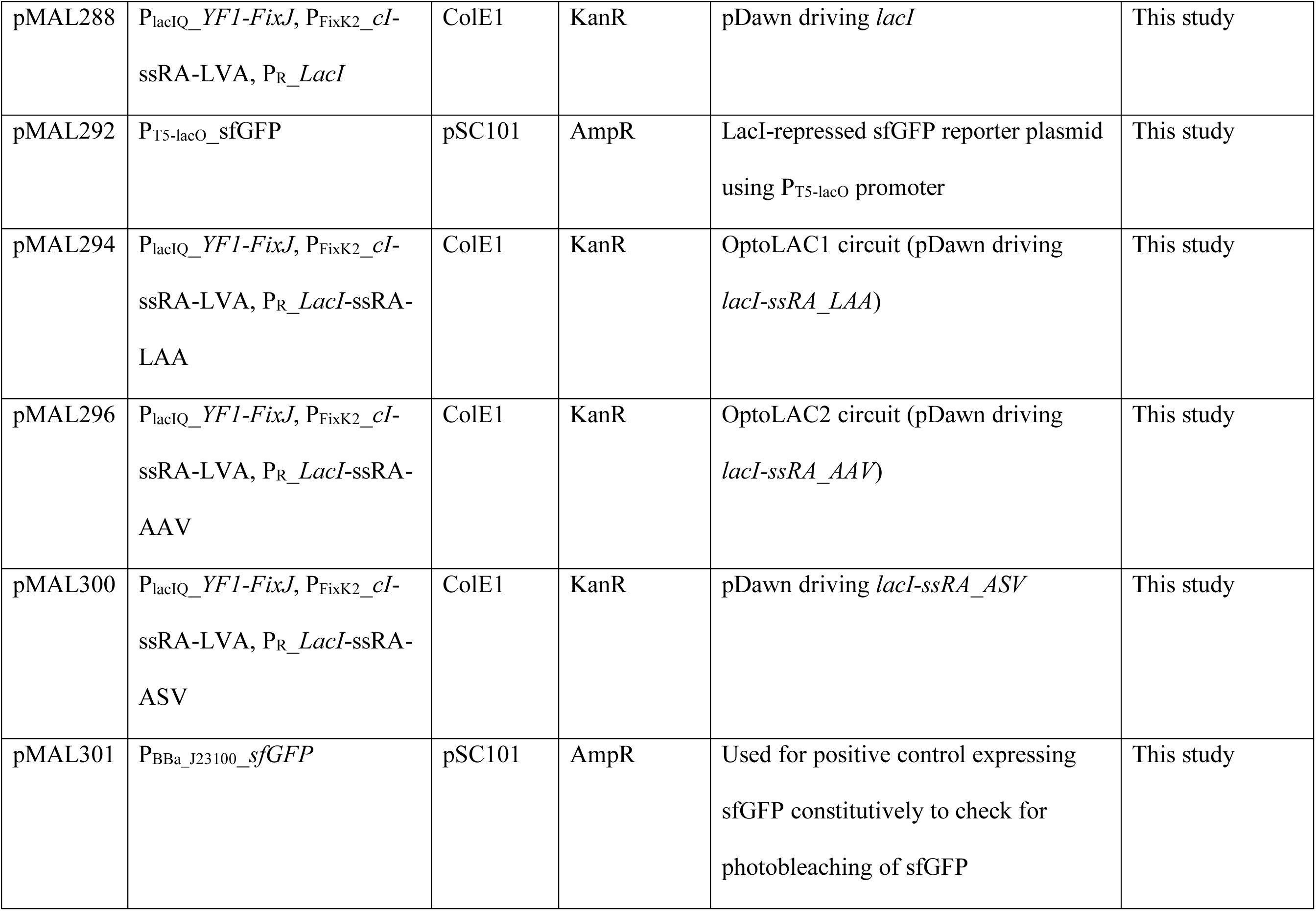
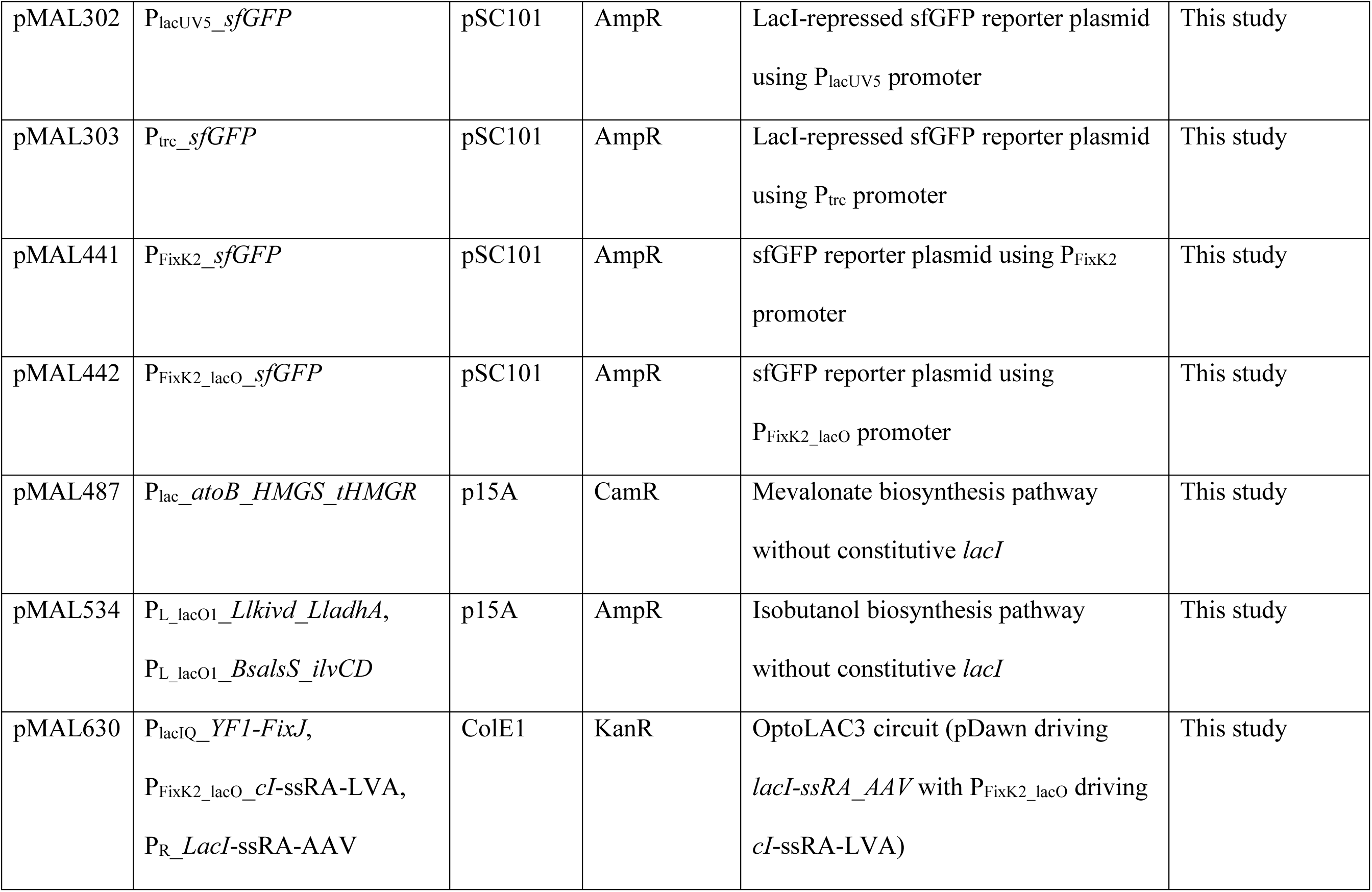
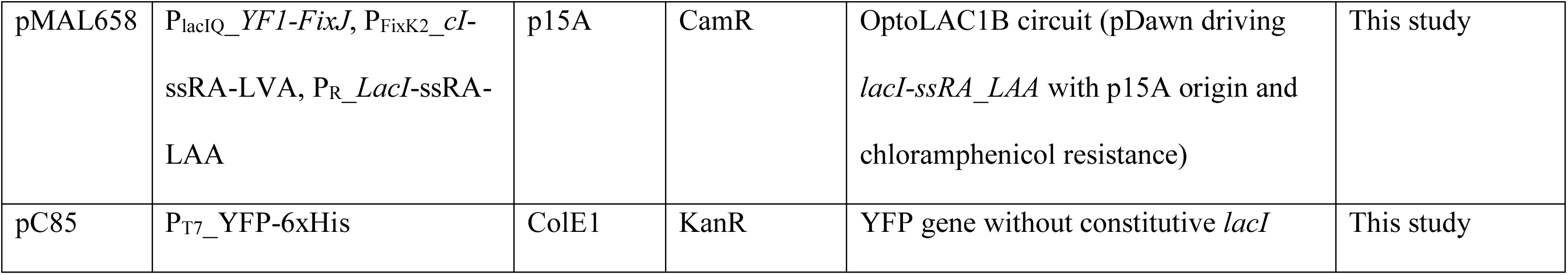
Plasmids used in this study

| Plasmid | Description | Origin of replication | Resistance marker | Additional information | Source |
| --- | --- | --- | --- | --- | --- |
| pDusk | P<sub>lacIQ</sub>_YF1-FixJ,<br>P<sub>FixK2</sub>_MCS | ColE1 | KanR | Template plasmid for dark-inducible gene expression | Ohlendorf et al.<sup>15</sup> |
| pDawn | P<sub>lacIQ</sub>_YF1-FixJ, P<sub>FixK2</sub>_cI-ssRA-LVA, P<sub>R</sub>_MCS | ColE1 | KanR | Template plasmid for blue light-inducible gene expression | Ohlendorf et al.<sup>15</sup> |
| pKD4 | FRT_*KanR*_FRT | R6K $\gamma$ | AmpR,<br>KanR | Used as a template for gene deletions | Datsenko and Wanner<sup>65</sup> |
| pKD46 | *araC*,<br>P<sub>araBAD</sub>_Gam_Beta_Exo | pSC101<br>(temp. sensitive) | AmpR | Used to induce gene deletion | Datsenko and Wanner<sup>65</sup> |
| pCP20 | *FLP*<sup>+</sup>, $\lambda$ cI857<sup>+</sup>, $\lambda$ P<sub>R</sub> | pSC101<br>(temp.<br>sensitive) | AmpR | Used for antibiotic marker recycling | Cherepanov and Wackernagel<sup>68</sup> |
| pET28a | P<sub>lacIQ</sub>_ *lacI*, P<sub>T7</sub>_MCS-6xHis | ColE1 | KanR | Contains constitutively expressed *lacI*; used for IPTG-inducible controls | Millipore Sigma |
| pMevT | P<sub>lac</sub>_ *atoB_HMGS_tHMGR* | p15A | CamR | Template plasmid for mevalonate production | Martin et al.<sup>24</sup> |
| pSA65 | P<sub>L_lacO1</sub>_ *Llkivd_LladhA* | ColE1 | AmpR | Template plasmid for isobutanol production | Atsumi et al.<sup>63</sup> |
| pSA69 | P<sub>L_lacO1</sub>_ *BsalsS_ilvCD* | p15A | KanR | Template plasmid for isobutanol production | Atsumi et al.<sup>27</sup> |
| pCri-8b | P<sub>T7</sub>_YFP-6xHis, P<sub>lacIQ</sub>_ *lacI* | ColE1 | KanR | IPTG-inducible YFP with constitutively expressed *lacI* | Goulas et al.<sup>30</sup> |
| pMAL207 | Empty vector | pSC101 | AmpR | Used for negative control to subtract autofluorescence | This study |
| pMAL288 | P<sub>lacIQ</sub>_YFI-FixJ, P<sub>FixK2</sub>_cI-ssRA-LVA, P<sub>R</sub>_LacI | ColE1 | KanR | pDawn driving *lacI* | This study |
| pMAL292 | P<sub>T5-lacO</sub>_sfGFP | pSC101 | AmpR | LacI-repressed sfGFP reporter plasmid using P<sub>T5-lacO</sub> promoter | This study |
| pMAL294 | P<sub>lacIQ</sub>_YFI-FixJ, P<sub>FixK2</sub>_cI-ssRA-LVA, P<sub>R</sub>_LacI-ssRA-LAA | ColE1 | KanR | OptoLAC1 circuit (pDawn driving *lacI-ssRA_LAA*) | This study |
| pMAL296 | P<sub>lacIQ</sub>_YFI-FixJ, P<sub>FixK2</sub>_cI-ssRA-LVA, P<sub>R</sub>_LacI-ssRA-AAV | ColE1 | KanR | OptoLAC2 circuit (pDawn driving *lacI-ssRA_AAV*) | This study |
| pMAL300 | P<sub>lacIQ</sub>_YFI-FixJ, P<sub>FixK2</sub>_cI-ssRA-LVA, P<sub>R</sub>_LacI-ssRA-ASV | ColE1 | KanR | pDawn driving *lacI-ssRA_ASV* | This study |
| pMAL301 | P<sub>BBa_J23100</sub>_sfGFP | pSC101 | AmpR | Used for positive control expressing sfGFP constitutively to check for photobleaching of sfGFP | This study |
| pMAL302 | P<sub>lacUV5</sub>*_sfGFP* | pSC101 | AmpR | LacI-repressed sfGFP reporter plasmid using P<sub>lacUV5</sub> promoter | This study |
| pMAL303 | P<sub>trc</sub>*_sfGFP* | pSC101 | AmpR | LacI-repressed sfGFP reporter plasmid using P<sub>trc</sub> promoter | This study |
| pMAL441 | P<sub>FixK2</sub>*_sfGFP* | pSC101 | AmpR | sfGFP reporter plasmid using P<sub>FixK2</sub> promoter | This study |
| pMAL442 | P<sub>FixK2_lacO</sub>*_sfGFP* | pSC101 | AmpR | sfGFP reporter plasmid using P<sub>FixK2_lacO</sub> promoter | This study |
| pMAL487 | P<sub>lac</sub>*_atoB_HMGS_tHMGR* | p15A | CamR | Mevalonate biosynthesis pathway without constitutive *lacI* | This study |
| pMAL534 | P<sub>L_lacO1</sub>*_Llkivd_LladhA*,<br>P<sub>L_lacO1</sub>*_BsalsS_ilvCD* | p15A | AmpR | Isobutanol biosynthesis pathway without constitutive *lacI* | This study |
| pMAL630 | P<sub>lacIQ</sub>*_YFI-FixJ*,<br>P<sub>FixK2_lacO</sub>*_cI-ssRA-LVA*,<br>P<sub>R_LacI</sub>*-ssRA-AAV* | ColE1 | KanR | OptoLAC3 circuit (pDawn driving *lacI-ssRA_AAV* with P<sub>FixK2_lacO</sub> driving *cI-ssRA-LVA*) | This study |
| pMAL658 | P<sub>lacIQ</sub>_YFI-FixJ, P<sub>FixK2</sub>_cI-ssRA-LVA, P<sub>R</sub>_LacI-ssRA-LAA | p15A | CamR | OptoLAC1B circuit (pDawn driving *lacI-ssRA_LAA* with p15A origin and chloramphenicol resistance) | This study |
| pC85 | P<sub>T7</sub>_YFP-6xHis | ColE1 | KanR | YFP gene without constitutive *lacI* | This study |

**Supplementary Table 2:** Bacterial strains used in this study

| Strain | Genotype | Plasmid(s) | Additional information | Source |
| --- | --- | --- | --- | --- |
| MG1655 | str. K F- λ- rph-1 | None | *E. coli* K strain | <sup>70</sup> |
| BL21 DE3 | str. B F- ompT gal dcm lon hsdSB(rB-mB-) $\lambda$(DE3 [lacI lacUV5-T7p07 ind1 sam7 nin5]) [malB+]K-12(λS) | None | *E. coli* B strain containing $\lambda$ prophage | <sup>71</sup> |
| MG1655 $\Delta lacI::KanR$ | F- λ- rph-1<br>$\Delta lacI::FRT$-KanR-FRT | None | MG1655 containing *lacI* deletion | Orman and<br>Brynildsen<sup>17</sup> |
| EMAL52 | MG1655 $\Delta lacI::FRT$ | None | OptoMG: Base K strain for chemical<br>production | This study |
| EMAL57 | MG1655 $\Delta lacI::FRT$ | pMAL288,<br>pMAL292 | OptoMG containing pDawn_*lacI* and<br>sfGFP driven by P<sub>T5-lacO</sub> | This study |
| EMAL68 | MG1655 $\Delta lacI::FRT$ | pMAL294,<br>pMAL292 | OptoMG containing OptoLAC1 and sfGFP<br>driven by P<sub>T5-lacO</sub> | This study |
| EMAL69 | MG1655 $\Delta lacI::FRT$ | pMAL296,<br>pMAL292 | OptoMG containing OptoLAC2 and sfGFP<br>driven by P<sub>T5-lacO</sub> | This study |
| EMAL71 | MG1655 $\Delta lacI::FRT$ | pMAL300,<br>pMAL292 | OptoMG containing pDawn_*lacI*-<br>*ssRA_ASV* and sfGFP driven by P<sub>T5-lacO</sub> | This study |
| EMAL77 | MG1655 $\Delta lacI::FRT$ | pET28a, pMAL292 | OptoMG containing IPTG-induced control<br>for sfGFP production | This study |
| EMAL135 | MG1655 $\Delta lacI::FRT$ | pET28a, pMAL487 | OptoMG containing IPTG-induced control for mevalonate production | This study |
| EMAL152 | MG1655 $\Delta lacI::FRT$ | pDusk, pMAL441 | OptoMG containing YF1-FixJ and sfGFP driven by $P_{FixK2}$ | This study |
| EMAL153 | MG1655 $\Delta lacI::FRT$ | pDusk, pMAL442 | OptoMG containing YF1-FixJ and sfGFP driven by $P_{FixK2\_lacO}$ | This study |
| EMAL199 | MG1655 $\Delta lacI::FRT$ | pMAL294,<br>pMAL534 | OptoMG containing OptoLAC1 and isobutanol production pathway | This study |
| EMAL200 | MG1655 $\Delta lacI::FRT$ | pMAL296,<br>pMAL534 | OptoMG containing OptoLAC2 and isobutanol production pathway | This study |
| EMAL201 | MG1655 $\Delta lacI::FRT$ | pET28a, pMAL534 | OptoMG containing IPTG-induced control for isobutanol production | This study |
| EMAL208 | MG1655 $\Delta lacI::FRT$ | pMAL294,<br>pMAL487 | OptoMG containing OptoLAC1 and mevalonate production pathway | This study |
| EMAL209 | MG1655 $\Delta lacI::FRT$ | pMAL296,<br>pMAL487 | OptoMG containing OptoLAC2 and mevalonate production pathway | This study |
| EMAL224 | BL21 DE3<br>*ΔlacI::FRT* | None | Precursor to EMAL255: BL21 DE3 with single deletion of *lacI* | This study |
| EMAL229 | MG1655 *ΔlacI::FRT* | pMAL630,<br>pMAL207 | Negative control OptoMG strain for subtraction of autofluorescence | This study |
| EMAL230 | MG1655 *ΔlacI::FRT* | pMAL630,<br>pMAL292 | OptoMG containing OptoLAC3 and sfGFP driven by P<sub>T5-lacO</sub> | This study |
| EMAL231 | MG1655 *ΔlacI::FRT* | pMAL630,<br>pMAL301 | Positive control OptoMG strain containing sfGFP driven constitutively by P<sub>BBa_J23100</sub> | This study |
| EMAL235 | MG1655 *ΔlacI::FRT* | pMAL630,<br>pMAL487 | OptoMG containing OptoLAC3 and mevalonate production pathway | This study |
| EMAL239 | MG1655 *ΔlacI::FRT* | pMAL630,<br>pMAL534 | OptoMG containing OptoLAC3 and isobutanol production pathway | This study |
| EMAL255 | BL21 DE3 *ΔlacI*<br>*ΔlacI (DE3)::SpecR* | None | OptoBL: Base B strain for chemical production | This study |
| EMAL267 | MG1655 *ΔlacI::FRT* | pMAL294,<br>pMAL302 | OptoMG containing OptoLAC1 and sfGFP driven by P<sub>lacUV5</sub> | This study |
| EMAL268 | MG1655 $\Delta lacI::FRT$ | pMAL294,<br>pMAL304 | OptoMG containing OptoLAC1 and sfGFP<br>driven by $P_{trc}$ | This study |
| EMAL283 | BL21 DE3 | pCri-8b | IPTG-induced OptoBL control strain for<br>YFP driven by $P_{T7}$ | This study |
| EMAL284 | BL21 DE3 $\Delta lacI$<br>$\Delta lacI$ (DE3)::SpecR | pC85, pMAL658 | OptoBL containing OptoLAC1B and YFP<br>driven by $P_{T7}$ | This study |

Red highlight indicates RBS^69^; yellow highlight indicates *lacO* sequence.

**Supplementary Figure 1.**
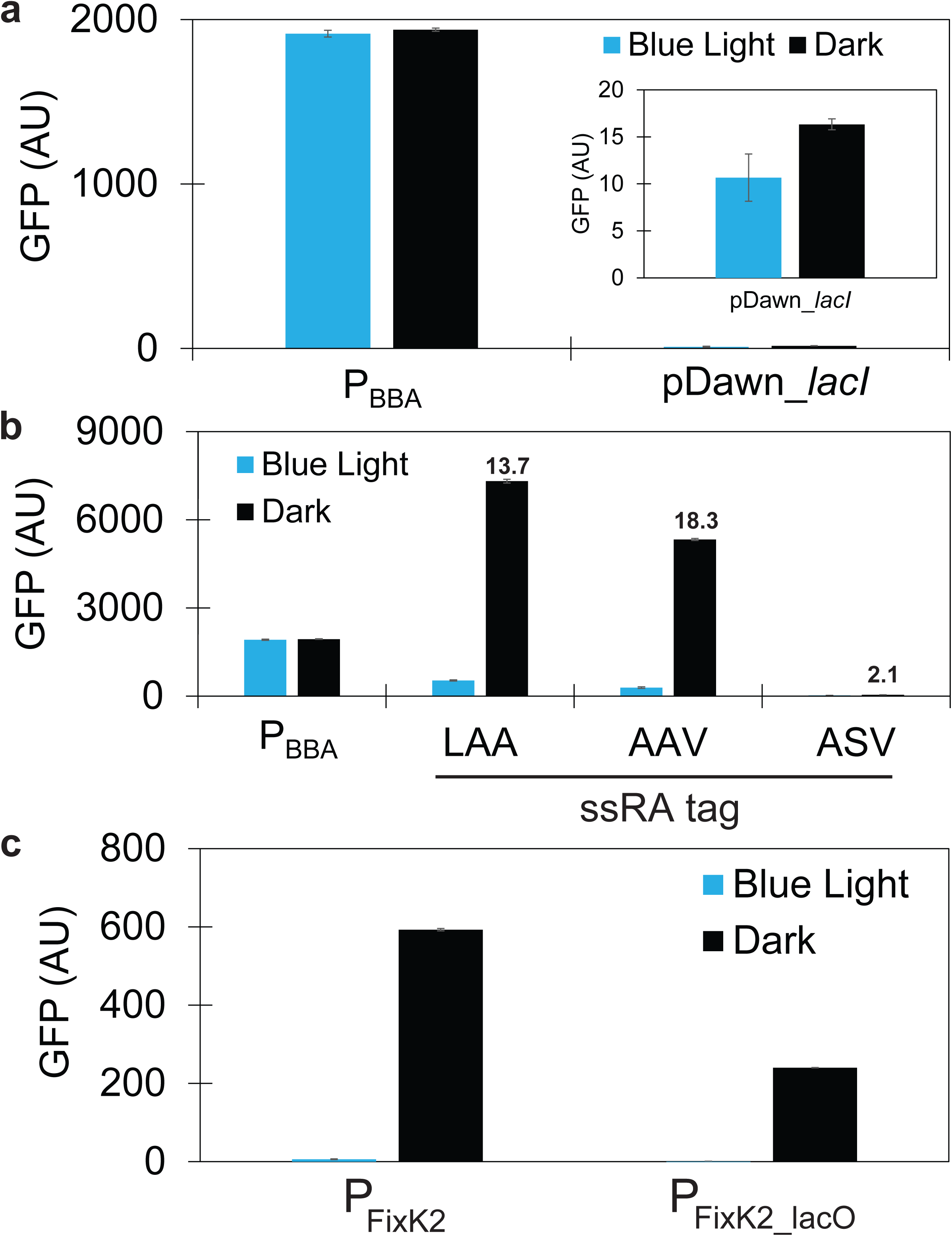
Development of OptoLAC circuits. **(a)** GFP expression in blue light or darkness from the constitutive P_BBA_ promoter (EMAL231) or from P_T5-lacO_ in a strain where *lacI* expression is controlled by pDawn (EMAL57). The inbox rescales the Y-axis to show marginal levels of pDawn-*lacI* darkness induction. (b) GFP expression in blue light or darkness from P_BBA_ (EMAL231) or from P_T5-lacO_ in strains containing the pDawn system controlling LacI fused to ssRA tags terminating in LAA (EMAL68, which gave rise to OptoLAC1), AAV (EMAL69, which gave rise to OptoLAC2), or ASV (EMAL71). (c) Effect of inserting a *lacO* site on the strength of the P_FixK2_ promoter, which is used in OptoLAC3. GFP expression driven by the native P_FixK2_ (pMAL441) or P_FixK2-lacO_ (pMAL442) promoter using the pDusk system (EMAL152 and EMAL153, respectively). All data shown as median values of 10,000 single-cell flow cytometry events; error bars represent one standard deviation of replicates exposed to the same conditions (n = 3 biologically independent 150 µL samples). All experiments were repeated at least three times.

**Supplementary Figure 2.**
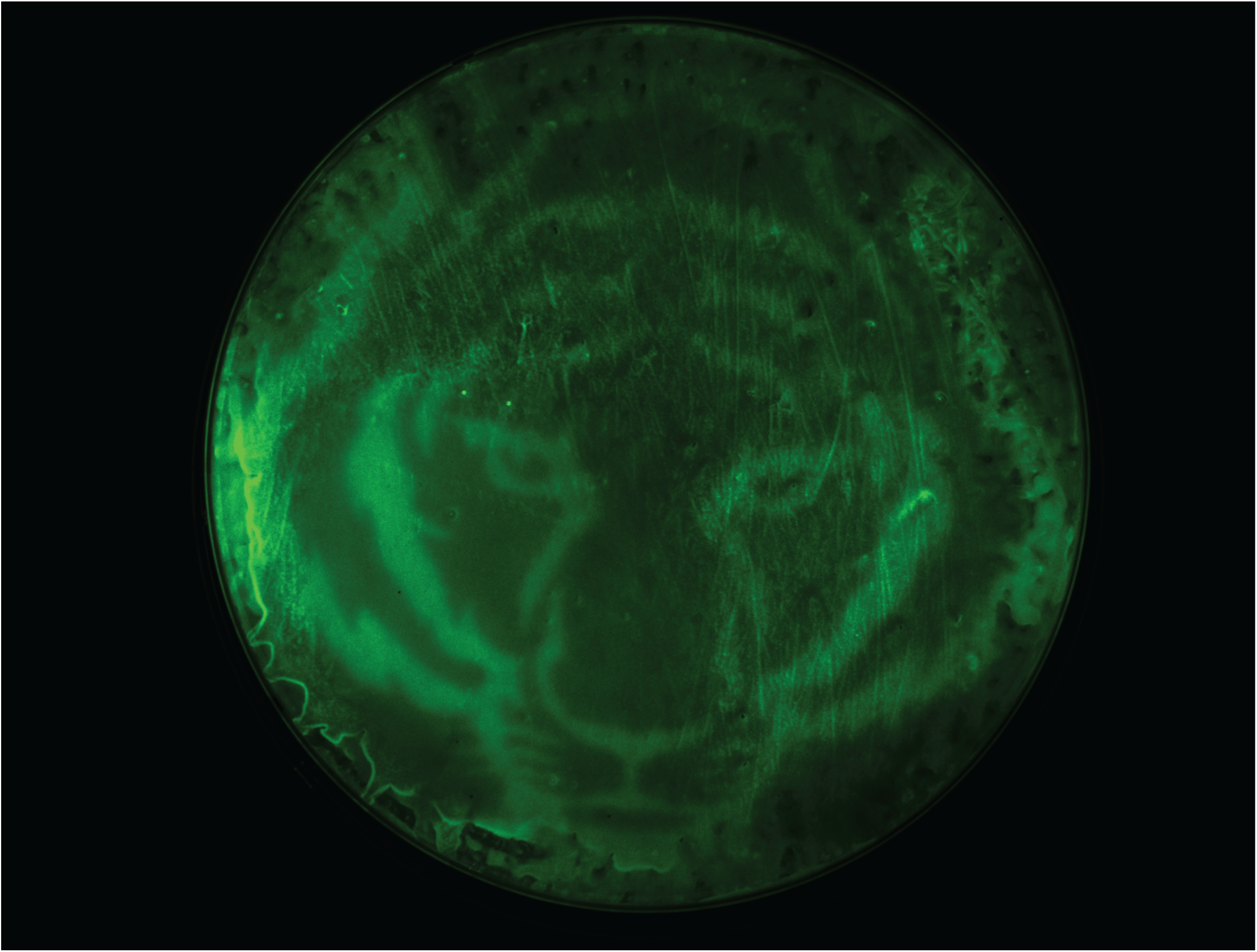
Spatial control of GFP expression on an LB agar plate. LB + Kan + Amp agar plate containing a lawn of EMAL68 (OptoLAC1 driving GFP) illuminated with a projection of a tiger image for 16 hours.

**Supplementary Figure 3.**
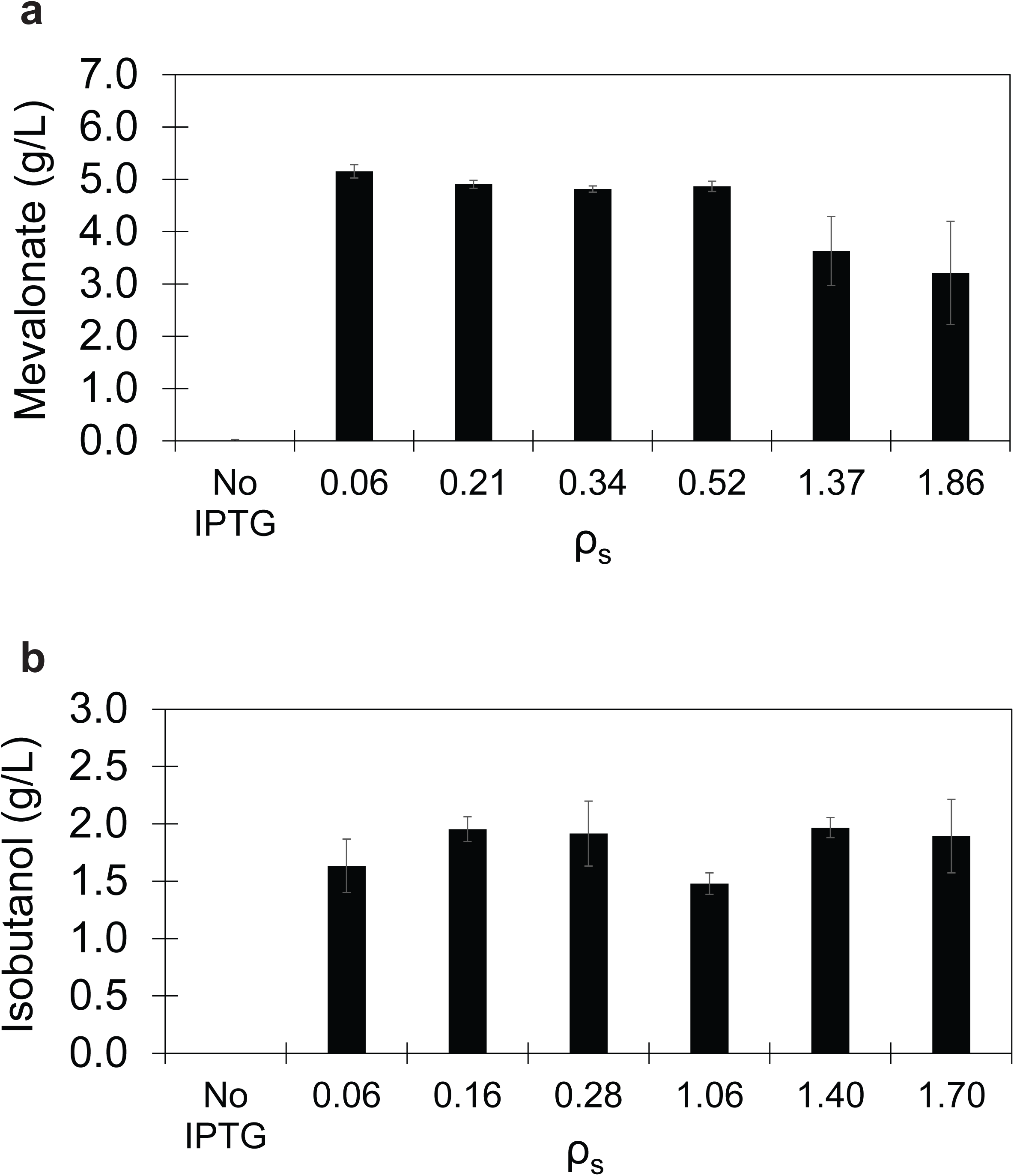
Optimization of chemical production with IPTG induction. **(a)** Mevalonate production from pMAL487 by EMAL135 when induced with 1mM IPTG at different cell densities. **(b)** Isobutanol production from pMAL534 by EMAL201 when induced with 1mM IPTG at different cell densities. All data are shown as mean values; error bars represent the standard deviation of at least three biologically independent 1-mL sample replicates exposed to the same conditions. All experiments were repeated at least three times.

